# The Notch Ligand Jagged1 is Required for the Formation, Maintenance, and Survival of Hensen Cells in the Mouse Cochlea

**DOI:** 10.1101/2020.05.04.076448

**Authors:** Elena Chrysostomou, Luyi Zhou, Yuanzhao L. Darcy, Kaley A. Graves, Angelika Doetzlhofer, Brandon C. Cox

## Abstract

During cochlear development, the Notch ligand JAGGED 1 (JAG1) plays an important role in the specification of the prosensory region, which gives rise to sound-sensing hair cells and neighboring supporting cells (SCs). While JAG1’s expression is maintained in SCs through adulthood, the function of JAG1 in SC development is unknown. Here, we demonstrate that JAG1 is essential for the formation and maintenance of Hensen cells (HeCs), a highly specialized SC-subtype located at the edge of the auditory epithelium. Deletion of *Jag1* at the onset of differentiation, at stage E14.5, disrupted HeC formation. Similar loss of HeCs was observed when *Jag1* was deleted at P0/P1 and fate-mapping analysis revealed that in the absence of *Jag1* some HeCs die, but others convert into neighboring Claudius cells. In support of a role for JAG1 in cell survival, genes involved in mitochondrial function and protein synthesis were downregulated in P0 cochlea lacking *Jag1*.

## INTRODUCTION

The canonical Notch signaling cascade plays multiple roles in the development of sensory structures in the vertebrate inner ear. In this pathway, activation of the Notch receptor by membrane-bound ligands located on neighboring cells leads to the release of the Notch intercellular domain, which as part of a larger transcription complex, activates the transcription downstream target genes (Kopan and Ilagan, 2009).

During inner ear sensory development, Notch signaling operates through two distinct modes of signaling, lateral inhibition and lateral induction (Eddison et al., 2000, Daudet and Zak, 2020). Notch-mediated lateral induction first defines and maintains the prosensory domains that give rise to highly specialized sensory epithelia composed of hair cells (HCs) and surrounding supporting cells (SCs). The Notch ligand JAGGED1 (JAG1) and the downstream transcription factor SOX2 are essential for the early role of Notch signaling in prosensory progenitor specification and maintenance (Daudet and Lewis, 2005, Hartman et al., 2010, Pan et al., 2010, Neves et al., 2011) and early loss of JAG1 abolishes or greatly reduces the pool of inner ear prosensory progenitors (Brooker et al., 2006, Kiernan et al., 2006, Petrovic et al., 2014).

As prosensory progenitor cells differentiate into HCs and SCs during late embryogenesis, Notch signaling plays a key role in limiting the number of HCs that form in a process termed lateral inhibition. Activated by ATOH1, a transcriptional activator and master regulator of HC formation (Bermingham et al., 1999, Chen et al., 2002, Woods et al., 2004), nascent HCs express the Notch ligands *Delta-like 1* (*Dll1*), *Delta-like 3* (*Dll3*), and *Jagged 2* (*Jag2*) (Adam et al., 1998, Lanford et al., 1999, Morrison et al., 1999, Hartman et al., 2007). Activation of Notch 1 receptor signaling by DLL1 and JAG2 in adjacent prosensory progenitor cells inhibits these cells from becoming HCs (Lanford et al., 1999, Kiernan et al., 2005, Brooker et al., 2006).

Important effectors of this HC-repressive function mediated by Notch signaling are members of the HES/HEY family. HES/HEY proteins are transcriptional repressors, which antagonize HC formation by repressing *Atoh1* expression and ATOH1 activity (Zine et al., 2001, Zheng and Gao, 2000, Li et al., 2008, Doetzlhofer et al., 2009, Tateya et al., 2011).

We recently uncovered that canonical Notch signaling, in addition to its role in HC fate repression, is required for the differentiation and survival of cochlear SCs (Campbell et al., 2016). However, the Notch ligand(s) and receptor(s) involved in this process are unknown. A potential candidate for instructing SC development is the Notch ligand JAG1. JAG1, which initially is expressed by cochlear and vestibular prosensory progenitors, continues to be highly expressed in SCs (Morrison et al., 1999). While other Notch ligands are down-regulated during the first postnatal week, coinciding with the early phase of cochlear HC and SC maturation, SC-specific JAG1 expression continues throughout adulthood where its function is unknown (Maass et al., 2015, Murata et al., 2006, Hartman et al., 2007, Oesterle et al., 2008).

In the present study, we investigated the role of JAG1 in differentiating and maturing SCs. Our analysis revealed that deletion of *Jag1* at the onset of cochlear differentiation, alters the patterning and cellular morphology of SCs and leads to a down-regulation of genes involved in mitochondrial function and protein synthesis within the sensory epithelium. Furthermore, using morphological analyses and fate-mapping, we show that *Jag1* deletion prior to and after cochlear differentiation resulted in loss of a specialized SC subtype called Hensen cells (HeCs). Functional analysis of adult *Jag1* mutant animals in which *Jag1* was deleted at the perinatal stage revealed mild hearing deficits at low frequencies. Together, our results suggest that *Jag1*-mediated Notch signaling in cochlear SCs is critical for the formation, maintenance, and survival of HeCs.

## RESULTS

The mammalian auditory sensory epithelium houses several distinct SC subtypes including HeCs, Deiters’ cells (DCs), inner and outer pillar cells (IPCs, OPCs), inner phalangeal cells (IPhCs), and border cells (BCs) (Fig. 1A). To investigate the function of JAG1 in differentiating SCs, we made use of *Jag1^loxP/loxP^* (*Jag1^fx/fx^*) mice, previously established by Dr. Julian Lewis’s group (Brooker et al., 2006) and *Sox2*^*CreERT2/+*^ mice (Arnold et al., 2011). In the cochlea, *Sox2^CreERT2/+^* is highly expressed in HC and SC precursors and by one week of age its expression becomes restricted to terminal differentiated SCs (Walters et al., 2015, Gu et al., 2016). We timed-mated *Sox2^CreERT2/+^::Jag1^fx/fx^* with *Jag1^fx/fx^* mice and administered tamoxifen to pregnant dams at stage embryonic day (E) 14.5 (Fig. 1B), which bypasses the earlier requirement for JAG1 in cochlear prosensory progenitor specification and maintenance (Brooker et al., 2006, Kiernan et al., 2006, Hao et al., 2012, Campbell et al., 2016). To control for *Sox2* haploinsufficiency, we also collected animals that did not receive tamoxifen treatment (untreated). We harvested control (*Jag1^fx/fx^* treated, *Jag1^fx/fx^* untreated, and *Sox2^CreERT2/+^::Jag1^fx/fx^* untreated) and *Sox2^CreERT2/+^::Jag1^fx/fx^* treated [*Jag1* conditional knockout (CKO)] embryos at E18.5/postnatal day (P) 0.

**Figure 1.**
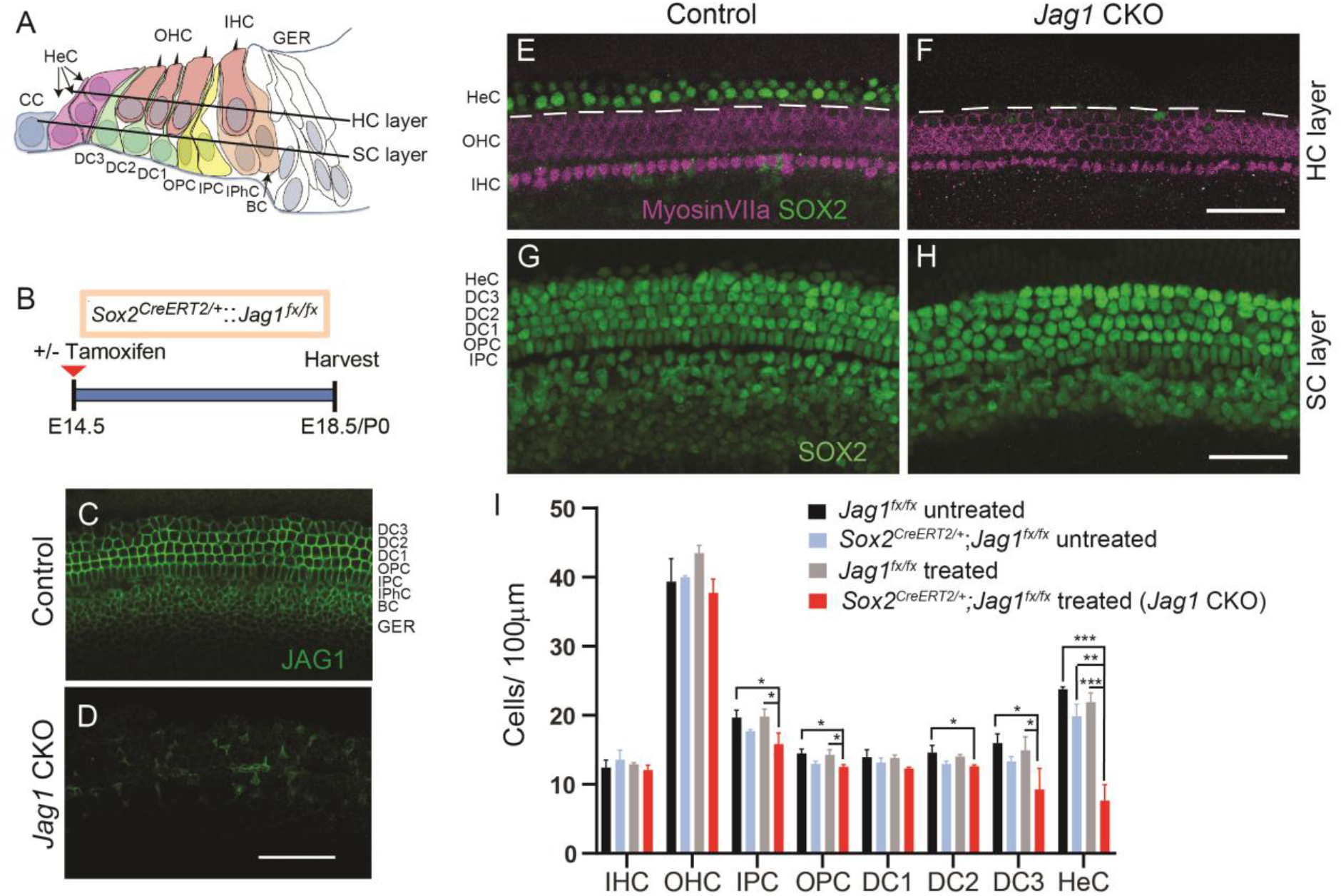
Deletion of *Jag1* from stage E14.5 progenitor cells disrupts HeC formation. **(A)** Schematic of the cellular composition of the neonatal auditory sensory epithelium. Abbreviations: BC, Border cell; CC, Claudius cell; DC1, Deiters’ cell row 1; DC2, Deiters’ cell row 2; DC3, Deiters’ cell row 3; GER, greater epithelial ridge; HeC, Hensen’s cell; HC, hair cell; IHC, inner hair cell; IPC, inner pillar cell; OHC, outer hair cell; OPC, outer pillar cell; SC, supporting cell. **(B)** Experimental strategy. Timed pregnant dams were injected with tamoxifen and progesterone at E14.5. *Jag1* conditional knockout (CKO) (*Sox2^CreERT2/+^::Jag1^fx/fx^* treated) mice and control (*Jag1^fx/fx^* treated) littermates were analyzed at E18.5/P0. SOX2 immuno-staining labels SC nuclei including HeCs, while myosin VIIa immuno-staining labels IHCs and OHCs. Representative confocal images of the cochlear mid-turn are shown. To control for *Sox2* haploinsufficiency, E18.5/P0 *Sox2^CreERT2/+^::Jag1^fx/fx^* mice and *Jag1^fx/fx^* littermates that did not receive tamoxifen and progesterone (untreated) were also analyzed. **(C-D)** Validation of *Jag1* deletion. SC nuclear layer, immuno-stained for JAG1 (green), in *Jag1* CKO (D) and control (C, treated, *Jag1^fx/fx^*) mice. **(E-F)** HC and HeC phenotype in P0 control (E, *Sox2^CreERT2/+^::Jag1^fx/fx^* untreated) and P0 *Jag1* CKO (F) mice. Note control mice (E) contain 1-2 rows of SOX2^+^ (green) HeCs lateral to the myosin VIIa^+^ (magenta) OHCs (dashed white line) (see schematic in A). **(G-H)** SC phenotype in E18.5 control (G, *Jag1^fx/fx^* treated) and E18.5 *Jag1* CKO (H) mice. Note that DC nuclei are enlarged and misaligned in *Jag1* CKO mice. **(I)** Quantification of HC and SC subtypes in control and *Jag1* CKO mice. Data expressed as mean ± SD (n=3/group; two-way ANOVA, with Bonferroni correction was used to calculate p-values, *p<0.05, **p ≤0.001, ***p≤0.0001). Scale bar: 50 μm.

First, we analyzed JAG1 expression in control and *Jag1* CKO cochlear tissue to validate Cre-mediated deletion of *Jag1*. In control samples, JAG1 protein was localized at the surface of SCs, except for HeCs (Fig. 1C). By contrast, the majority of SCs in *Jag1* CKO animals lacked JAG1 protein expression (Fig. 1D). Next, we analyzed whether *Jag1* deletion disrupted the formation or patterning of SCs and/or HCs. To label HCs, we used immuno-staining against myosin VIIa, which labels both inner HCs (IHCs) and outer HCs (OHCs) (Hasson et al., 1995). To visualize SCs, we immuno-stained against SOX2, which at E18.5/P0 is highly expressed in the nucleus of all SC subtypes, including HeCs, and is expressed at a lower level in the nucleus of IHCs and OHCs (Kempfle et al., 2016). We found that *Jag1* deletion had no effect on the number (density) of IHCs or OHCs (Fig. 1E, F, I). Our analysis of the SC phenotype, however revealed a significant reduction in the number of HeCs in *Jag1* CKO mice compared to all three controls (Fig. 1E-I). In cochlear tissue of control animals, SOX2^+^ HeCs were located between the 3^rd^ row of DCs and Claudius cells (CCs), with their nuclei residing in both the HC and SC layers, and with 2-3 HeCs sitting on top of each other (Fig. 1A, E, G). By contrast, *Jag1* CKO cochlear tissue contained either no or only a few scattered HeCs within the HC and SC layers (Fig. 1F, H).

In addition, DCs in *Jag1* CKO mice had enlarged nuclei compared to control mice, and their arrangement appeared to be disorganized, suggesting defects in DC differentiation (Fig. 1G, H). Decreased numbers of DCs in the 2^nd^ and 3^rd^ row (DC2 and DC3) was observed in *Jag1* CKO samples compared to the *Jag1^fx/fx^* treated and/or *Jag1^fx/fx^* untreated control groups, but not when compared to *Sox2^CreERT2/+^::Jag1^fx/fx^* untreated controls (Fig. 1I). A similar result was observed for IPCs and OPCs (Fig. 1I), suggesting that *Jag1* deficiency combined with *Sox2* haploinsufficiency negatively affects the differentiation of DCs and PCs. Unfortunately, we were unable to address how *Jag1* deficiency combined with *Sox2* haploinsufficiency may impact DC and PC maturation as conditional deletion of *Jag1* at E14.5 resulted in early postnatal lethality. In summary, our analysis demonstrates that JAG1’s function is essential for the formation of HeCs in the mammalian cochlea.

To gain insight into how JAG1 may influence HeC development, we characterized JAG1-mediated gene expression changes in the developing auditory sensory epithelium using whole genome micro-arrays. We administered tamoxifen to timed pregnant dams at stage E14.5 and harvested *Jag1* CKO (*Sox2^CreERT2/+^::Jag1^fx/fx^*) and control (*Jag1^fx/fx^*) mice at stage E18.5/P0 (Fig. 2A). We extracted RNA from enzymatically purified control and *Jag1* CKO cochlear sensory epithelia, and analyzed RNA abundancy using the GeneChip® Mouse Clariom D Arrays (Fig. 2A). Using a one-way ANOVA model we determined genes that were significantly up-(red) or down-regulated (blue) in control versus *Jag1* CKO cochlear epithelial cells. After removing pseudogenes, imprinted, misaligned and low expressing genes (average mean log2 (control, *Jag1* CKO) <5.5), a total of 227 genes were differentially expressed between control and *Jag1* CKO samples with 55 genes being significantly up-regulated (red dots, p-value ≤0.05, log2(FC) ≥3σ) and 172 genes being down-regulated (blue dots, p-value ≤0.05, log2(FC) ≤−3σ) (Fig. 2B). To identify SC-specific genes that are regulated by JAG1-Notch signaling, we utilized published data to generate a high confidence list of 200 SC-specific genes (Maass et al., 2016, Burns et al., 2015). Our analysis revealed that 41 out of these 200 SC-specific genes (21%) were differentially regulated by JAG1-Notch signaling, with 32 being downregulated and 9 being upregulated in *Jag1* CKO samples compared to control (Fig. 2 C) (Supplemental Table 1, 2 genes marked $). Furthermore, the list of down-regulated genes included SC-specific genes previously reported to be positively regulated by Notch signaling (*Agr3*, *F2rl1, Hey1, Igfbp3*, *Lrrtm1, Lsamp*, *Ntf3, Rab3b, Shc3 and Trh*) (Fig. 2B) (Supplemental Table 1, genes marked #) (Maass et al., 2016, Campbell et al., 2016). Conversely, the list of up-regulated genes included a small subset of genes previously shown to be negatively regulated by Notch signaling in response to Notch inhibition (*Cdh4*, *Dpysl2*, *Robo2* and *Cpa6*) (Maass et al., 2016) (Supplemental Table 2, genes marked #). However, deletion of *Jag1* in the differentiating cochlea failed to significantly increase the expression of *Atoh1* or other early HC-specific transcription factors (Supplemental Table 2). This is consistent with our finding that *Jag1* deletion at E14.5 does not result in ectopic HC formation (Fig. 1F, I). To uncover the biological processes associated with JAG1-regulated genes, we performed gene ontology (GO) enrichment analysis using Database for Annotation, Visualization and Integrated Discovery (DAVID). We found that the list of down-regulated genes (p-value ≤0.05, log2(FC) ≤−2σ) was enriched for terms linked to mitochondrial function (oxidative phosphorylation) and mitochondrial dysfunction (Alzheimer, Parkinson and Huntington), as well as protein synthesis (ribosome, intracellular ribonucleoprotein complex) (Fig. 2D, E).

**Figure 2.**
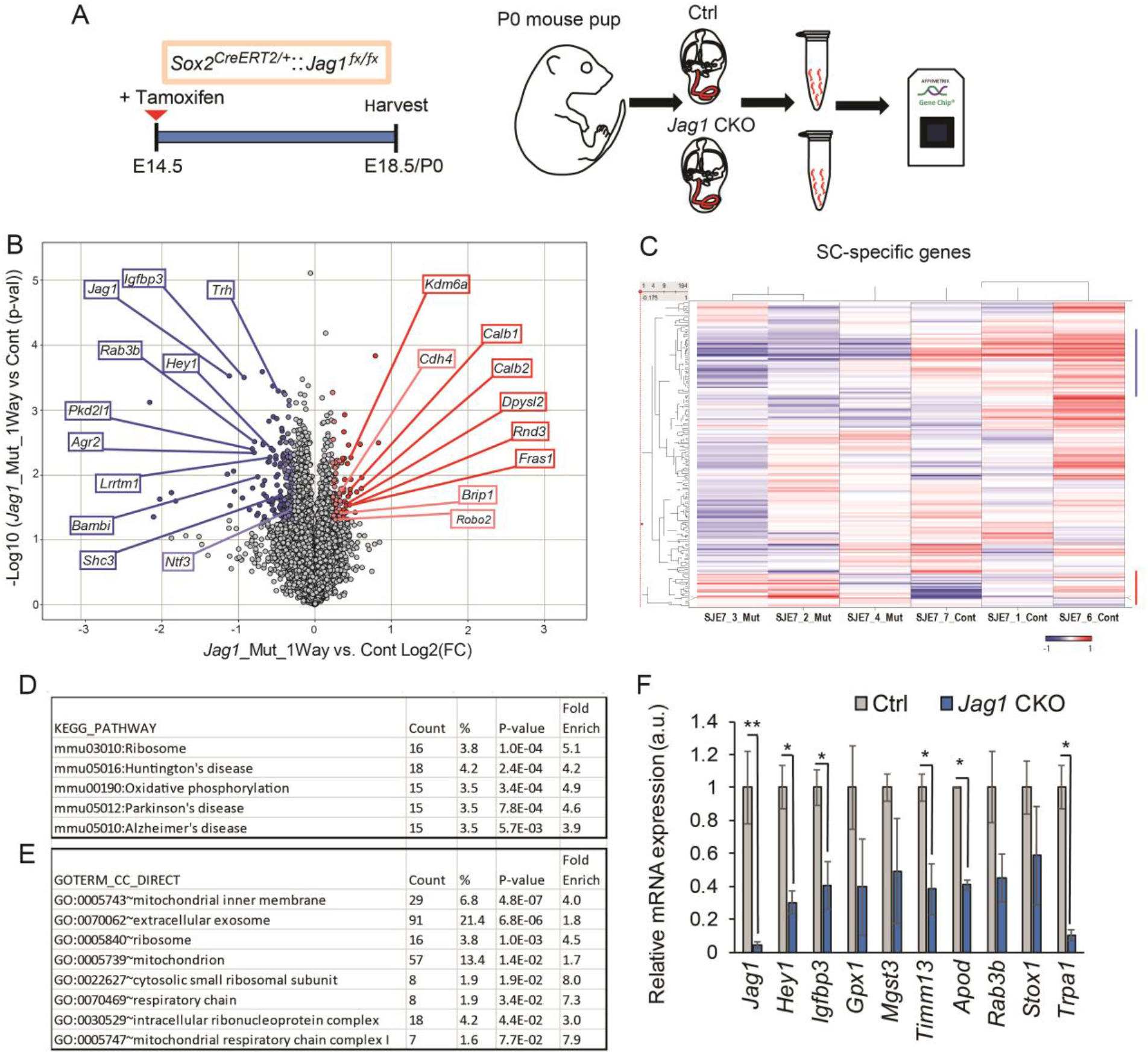
Loss of JAG1 leads to the downregulation of genes involved in mitochondrial function and protein synthesis. **(A)** Experimental strategy. Timed pregnant dams were injected with tamoxifen and progesterone at E14.5 and gene expression in cochlear epithelia from E18.5/P0 *Jag1* conditional knockout (CKO), (*Sox2^CreERT2/+^::Jag1^fx/fx^*) mice and control (Ctrl, *Jag1^fx/fx^*) littermates was analyzed using Clariom D mouse arrays. **(B)** Volcano plot presents the one-way ANOVA analysis of differential gene expression in *Jag1* CKO versus control samples (n=3 animals/group). Plotted as log2 fold-change (FC) (x-axis) versus −log10 p-value (y-axis). Dark blue (log2 (FC) <−6σ) and light blue (log2 (FC) <−3σ) dots represent downregulated genes. Dark red (log2 (FC) >6σ) and light red dots (log2 (FC) >3σ) represents upregulated genes. **(C)** Heat cluster map illustrating JAG1-mediated effects on a list of 200 SC-specific genes curated from published data. **(D-E)** Gene ontology (GO) analysis of genes down-regulated (p-value ≤0.05, log2 (FC) ≤2σ) in P0 *Jag1* CKO versus control (Ctrl) cochlear epithelia. Listed are significantly enriched (Bonferroni corrected p-value ≤0.05) pathways (D) and cellular component (E) GO terms. **(F)** RT-qPCR-based validation of genes of interest that were downregulated at P0 in *Jag1* CKO cochlear epithelia compared to Ctrl. Data expressed as mean ± SEM (minimum of n=3 animals/ group, two-tailed Students t-test was used to calculate p-values, *p<0.05, **p ≤0.001).

Next, we used qPCR to confirm *Jag1* deletion and validate a subset of SC-specific/enriched genes found to be down-regulated in *Jag1* mutant cochlear epithelia. The genes were selected based on their specific expression in HeCs (*Trpa1*) (Corey et al., 2004), and their association with biological processes of interest including oxidative stress and cell death [*Gpx1* (Ohlemiller et al., 2000), *Apod* (Ganfornina et al., 2008), *Mgst3* (Lu et al., 2016)], mitochondrial inner membrane transport (*Timm13*) (Rothbauer et al., 2001), and prosensory progenitor cell proliferation (*Stox1*) (Nie et al., 2015). We also included in our analysis a subset of genes, that have been reported to be positively regulated by Notch signaling in SC precursors (*Hey1, Igfbp3*) (Campbell et al., 2016) and in SCs (*Hey1*, *Igfbp3*, *Rab3b*) (Maass et al., 2016). We found that *Jag1*, *Hey1*, *Igfbp3*, *Timm13*, *Apod,* and *Trpa1* expression were significantly reduced in *Jag1* mutant cochlear epithelia. *Gpx1*, *Mgst3*, *Rab3b* and *Stox1* expression appeared reduced, but due to variability across samples the reduction was not significant. Taken together, our gene expression analysis confirms the observed defect in HeC formation and suggests a role for JAG1 in tissue homeostasis and metabolism.

In mice, HCs and SCs are not functional at birth and undergo a 2-week long process of postnatal maturation (Legendre et al., 2008, Lelli et al., 2009, Szarama et al., 2012). To address whether JAG1 function is required during postnatal maturation, we deleted *Jag1* after the completion of HC and SC differentiation. To that end, we injected *Jag1* CKO mice and their *Jag1^fx/fx^* littermates with 4-hydroxy-tamoxifen at P0/P1. To control for *Sox2* haploinsufficiency, we also collected *Sox2^CreERT2/+^::Jag1^fx/fx^* and *Jag1^fx/fx^* mice that did not receive 4-hydroxy-tamoxifen treatment (untreated) (Fig. 3A). Control (*Jag1^fx/fx^* treated, *Jag1^fx/fx^* untreated, and *Sox2^CreERT2/+^::Jag1^fx/fx^* untreated) and *Jag1* CKO mice were harvested and analyzed at P5 or P7. JAG1 immuno-staining confirmed the successful deletion of *Jag1* in cochlear tissue obtained from *Jag1* CKO mice (Fig. 3B, C). Next, we quantified HC and SC subtypes at stage P7 in *Jag1* CKO and control mice using the HC and SC markers myosin VIIa and SOX2 respectively. Quantification of SC numbers revealed a significant reduction in the number of HeCs in *Jag1* CKO mice compared to control mice (Fig. 3J). A recent report found that *Sox2* haploinsufficiency at the neonatal stage stimulates PC proliferation and the production of ectopic IHCs (Atkinson et al., 2018). Consistent with these recent findings, quantification of HC numbers in control and *Jag1* CKO mice uncovered a mild increase in the number of IHC in the two groups of mice that contained the *Sox2^CreERT2/+^* allele compared to the *Jag1^fx/fx^* treated and *Jag1^fx/fx^* untreated controls. Also we observed a mild, but significant increase in the number of IPCs in *Jag1* CKO mice compared to *Sox2^CreERT2/+^::Jag1^fx/fx^* untreated control animals (Fig 3J).

**Figure 3.**
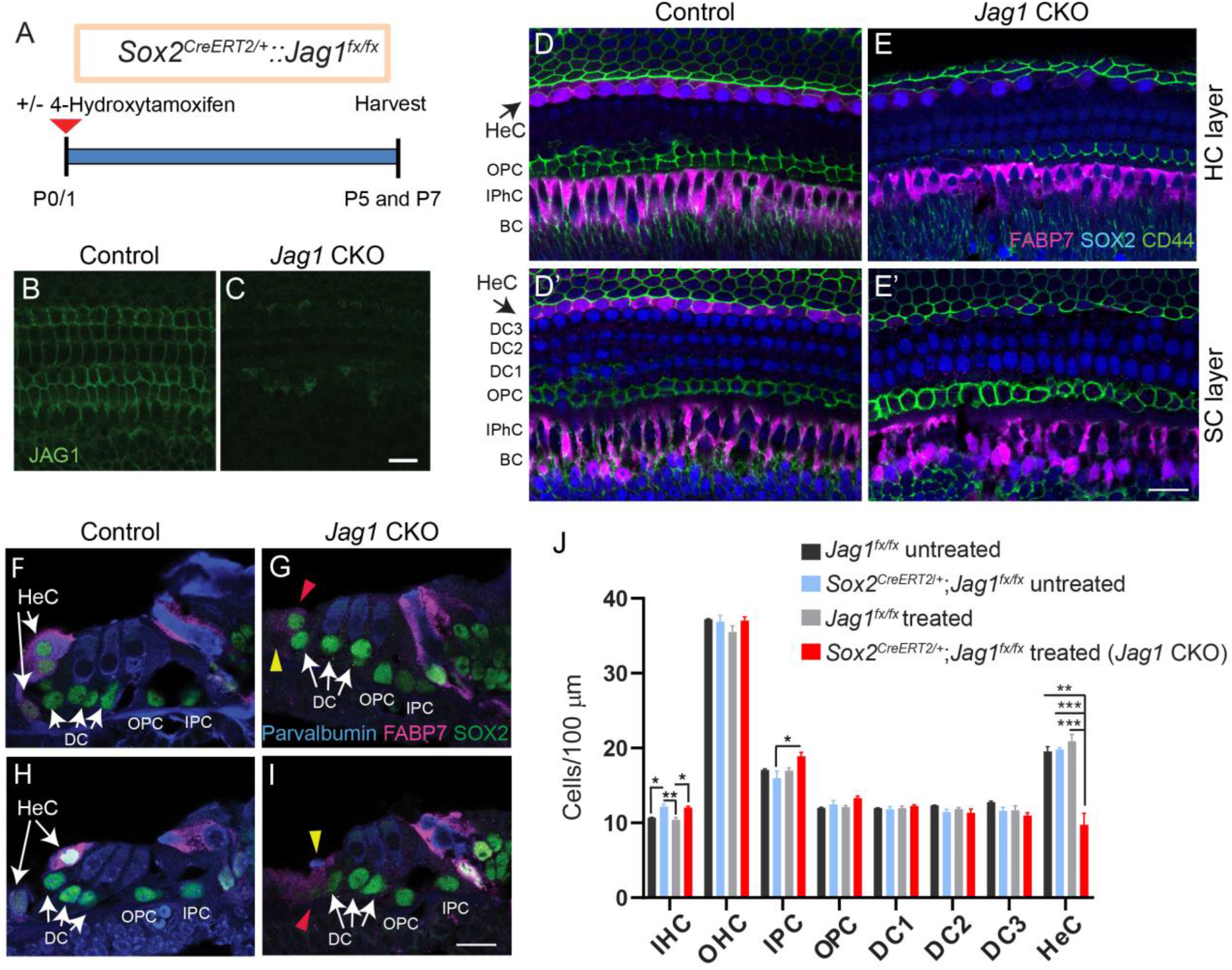
Deletion of *Jag1* from neonatal SCs results in the loss of HeCs. **(A)** Experimental strategy. *Sox2^CreERT2/+^::Jag1^fx/fx^* and *Jag1^fx/fx^* were treated with 4-hydroxy-tamoxifen at P0/P1 and their cochleae were analyzed at P5 or P7. Untreated control mice (*Sox2^CreERT2/+^::Jag1^fx/fx^* untreated and *Jag1^fx/fx^*, untreated) were also used. SOX2 immuno-staining labels SC nuclei including HeCs. FABP7 labels HeCs, IPhCs and BCs. Parvalbumin labels HCs and CD44 labels CCs and OPCs. Shown are representative single plane confocal images of the cochlear mid-turn. **(B-C)** Validation of conditional *Jag1* deletion. Shown is the SC nuclear layer, immuno-stained for JAG1 (green), in *Jag1* CKO (C) and control (B, *Jag1^fx/fx^* treated) mice. **(D-E’)** Top down view of the HC (D, E) and SC nuclear layers (D’, E’) in *Jag1* CKO (E, E’) and control (D, D’, *Sox2^CreERT2/+^::Jag1^fx/fx^* untreated) mice. Black arrows indicate the location of HeCs residing within the HC (D) and SC layer (D’) in control mice. **(F-I)** Cochlear cross-sections of *Jag1* CKO (G, I) and control mice (F, H, *Sox2^CreERT2/+^::Jag1^fx/fx^,* untreated). White arrows indicate 2-3 HeC nuclei that are stacked on top of each other in control mice. Red arrowheads indicate SOX2^+^ HeC-like cells (G, I) and yellow arrowheads indicate dying cell (I) and missing HeC (G) in *Jag1* CKO mice. **(J)** Quantification of HC and SC subtypes in control and *Jag1* CKO mice. Data expressed as mean ± SD (n=3/ group; two-way ANOVA, with Bonferroni correction was used to calculate p-values, *p<0.05, **p ≤0.001, ***p≤0.0001). Abbreviations: IHC, inner hair cell; OHC, outer hair cell; IPC, inner pillar cell; OPC, outer pillar cell; DC1, Deiters’ cell row 1; DC2, Deiters’ cell row 2; DC3, Deiters’ cell row 3; HeC, Hensen’s cell. Scale bar: 20 μm.

To confirm the observed HeC loss in *Jag1* CKO mice, we conducted additional experiments, in which we used a combination of fatty acid binding protein 7 (FABP7), SOX2, and CD44 immuno-staining to distinguish between HeCs, DCs, and CCs. In the postnatal cochlea, FABP7 is highly expressed in the cytoplasm of HeCs, IPhCs, and BCs (Saino-Saito et al., 2010), whereas CD44 is selectively expressed on the cell membranes of OPCs and CCs (Hertzano et al., 2010)(Fig. 3D, D’). In stage P5 control mice, 2-3 HeC nuclei (SOX2^+^, FABP7^+^) are stacked on top of each other (Fig. 3F, H), with a single row of HeCs within the SC nuclear layer and a row of stacked HeCs residing in the HC layer (Fig. 3D, D’). By contrast, the cochlea of *Jag1* CKO mice only contained a few scattered SOX2^+^ HeC-like nuclei within the HC layer and SC layer (Fig. 3E, E’). These HeC-like cells expressed FABP7 at a much-reduced level than HeCs in control mice and some SOX2^+^ HeC-like cells in *Jag1* CKO mice appeared to co-express CD44, suggesting that these cells may undergo a cell–fate conversion into CCs (Fig. 3E’).

To further investigate the HeC-like nuclei with CD44 expression, we performed fate-mapping experiments using *Sox2^CreERT2/+^::Jag1^fx/fx^::Rosa26^tdTomato/+^* and *Sox2^CreERT2/+^::Rosa26^tdTomato/+^* controls that were injected with tamoxifen at P0/P1 (Fig. 4A). In control samples collected at P7, all SOX2^+^ SC subtypes, including HeCs, expressed tdTomato (Tom, Fig. 4B, B”), and a small number of CD44^+^ CCs also expressed Tom (Fig. 4B’). Quantification of Tom^+^ cells lateral to the third row of DCs at P7 showed a decrease in the number of Tom^+^ cells and Tom^+^, SOX2^+^ cells in all turns of the cochlea when *Jag1* was deleted, which suggests that *Jag1* is important for the survival of some HeCs. In support of these findings, we frequently noted missing cells (Fig. 3G) as well as cellular debris (Fig. 3I) within the HeC region in *Jag1* CKO mice. However, staining with a marker of apoptosis, cleaved caspase-3, revealed no caspase3^+^ HeCs in the cochlea of *Jag1* CKO mice between P1-P4. There was also a ~2-fold increase in the number of cells labeled with both Tom and CD44 in *Jag1* CKO cochleae (Fig. 4C-F). Thus, some HeCs appear to lose SOX2 expression and convert into CCs.

**Figure 4.**
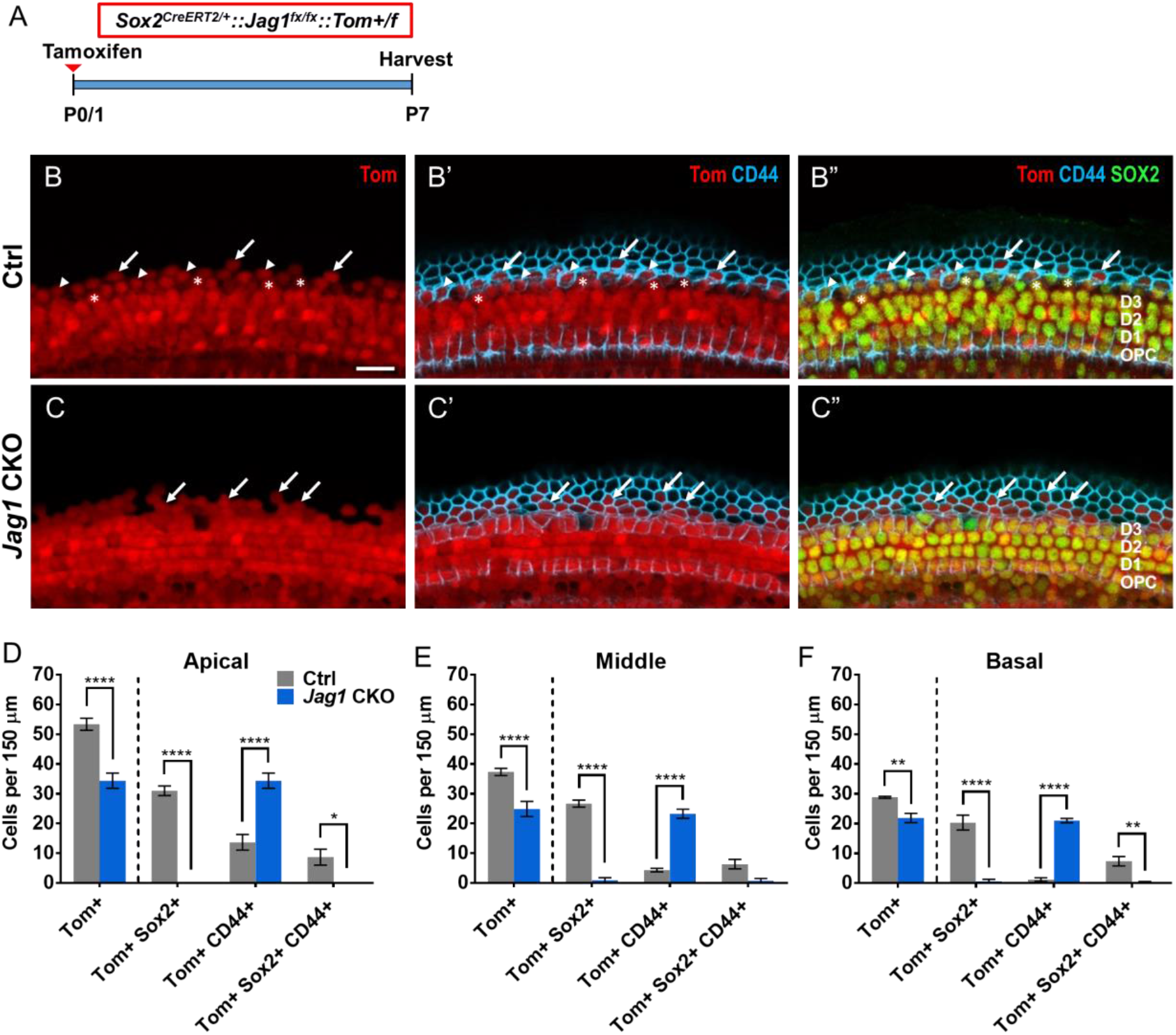
Fate-mapping analysis reveals that HeCs die or convert into CCs after deletion of *Jag1*. **(A)** Experimental strategy. *Sox2^CreERT2/+^::Jag1^fx/fx^::Rosa26^tdTomato/+^* (*Jag1* CKO) and *Sox2^CreERT2/+^::Rosa26^tdTomato/+^* (Ctrl) mice were injected with tamoxifen at P0/P1 and cochleae were analyzed at P7. **(B-C”)** Representative confocal images of the apical turn at P7 in Ctrl (B-B”) and *Jag1* CKO (C-C”) mice, showing expression of SOX2 (a SC marker, green), CD44 (a CC marker, blue) and tdTomato (Tom, red). Arrows indicate Tom^+^, CD44^+^ cells, * indicate Tom^+^, SOX2^+^ cells, and arrowheads indicate Tom^+^, SOX2^+^, CD44^+^ cells. **(D-F)** Quantification of Tom^+^ cells lateral to the 3^rd^ row of DCs that also express CD44 and/or SOX2 in the apical (D), middle (E), and basal (F) turns of the cochleae in Ctrl and *Jag1* CKO mice. Data are expressed as mean ± SEM. n=3-4/group; *p<0.05, **p<0.01, ****p<0.0001. Scale bar: 20 μm.

To investigate whether *Sox2* haploinsufficiency interacted with *Jag1* deletion to produce changes in HeC numbers, we used a different CreER line to delete *Jag1* from the neonatal cochlea*. Fgfr3-iCreER^T2/+^* is a transgenic allele that has been used extensively in cochlear studies and there have been no reports of haploinsufficiency (Atkinson et al., 2018, Kirjavainen et al., 2015). In contrast with *Sox2^CreERT2/+^* mice, *Fgfr3-iCreER^T2/+^* mice injected with tamoxifen at P0/P1 have CreER expression in ~100% of PCs and DCs, as well as in varying amounts of OHCs depending on the cochlear turn (Cox et al., 2012, McGovern et al., 2017). *Fgfr3-iCreER^T2/+^::Jag1^fx/fx^* and iCre-negative control mice were injected with tamoxifen at P0 and P1 to delete *Jag1*, which was confirmed by RT-PCR showing a second smaller band that lacks exon 4 in *Fgfr3-iCreER^T2/+^::Jag1^fx/fx^* mice (Fig. 5A-B). We quantified HCs and individual SC subtypes at P7 using myosin VIIa to label HCs and SOX2 to label SC nuclei (Fig. 5C-F’) and observed a significant loss of HeCs in *Fgfr3-iCreER^T2/+^::Jag1^fx/fx^* mice (Fig. 5I). To confirm that HeCs are missing, we employed two methods: 1) S100a1 immuno-staining to label the cytoplasm and nuclei of PCs and DCs (White et al., 2006, Buckiova and Syka, 2009) and 2) FABP7 immuno-staining to label the cytoplasm of HeCs (Saino-Saito et al., 2010). In control samples, there are two rows of SOX2^+^, FABP7^+^ cells located lateral to the S100a1^+^ third row of DCs (DC3) (Fig. 5E-E’, G-G’). However after *Jag1* deletion in *Fgfr3-iCreER^T2/+^::Jag1^fx/fx^* mice, there were no SOX2^+^ cells located lateral to DC3 cells (Fig. 5F-F’, H’) and there was a reduction in FABP7 expression (Fig. 5G, H). We also performed immuno-staining with CD44, to label CCs, which are located lateral to HeCs. In *Fgfr3-iCreER^T2/+^::Jag1^fx/fx^* mice, the CD44 expression pattern appeared similar to the control, however the CD44 staining was immediately adjacent to the SOX2^+^ DC3 nuclei (Fig. 5G”, H”). Together these results confirm that the missing SOX2^+^ cells are HeCs. In addition, there was a slight decrease in the number of OHCs, but no changes in the number of IHCs in *Fgfr3-iCreER^T2/+^::Jag1^fx/fx^* mice at P7 (Fig. 5C-D, I), which rules out the possibility of HeCs converting into HCs. Taken together these results indicate that *Sox2* haploinsufficiency is not responsible for the loss of HeCs after *Jag1* deletion.

**Figure 5.**
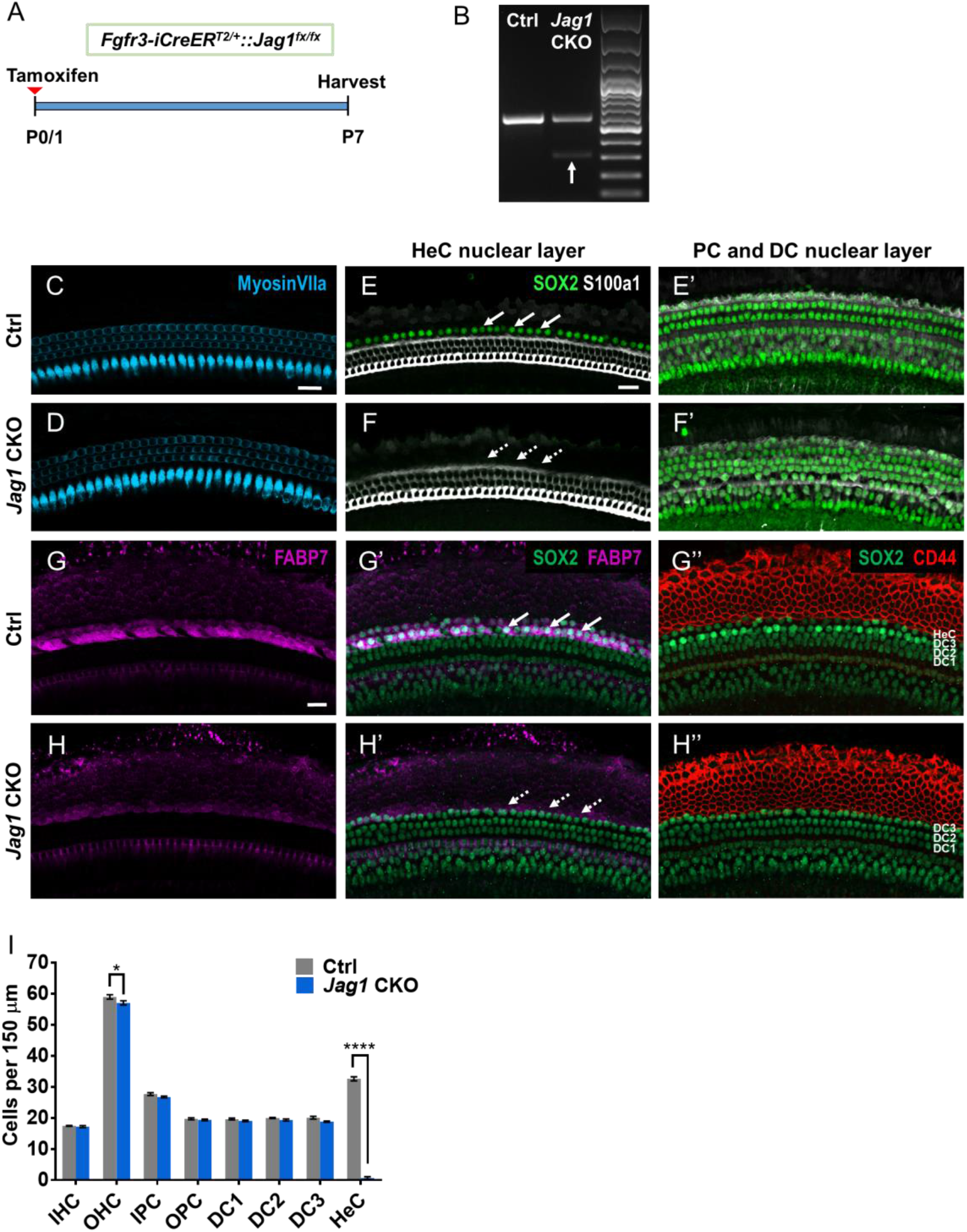
Massive loss of HeCs at P7 after neonatal deletion of *Jag1* from PCs and DCs. **(A)** Experimental strategy. *Fgfr3-iCreER^T2/+^*::*Jag1^fx/fx^* (*Jag1* CKO) and *Jag1^fx/fx^* (Ctrl) mice were injected with tamoxifen at P0/P1 and cochleae were analyzed at P7. **(B)** RT-PCR results using RNA extracted from P7 Ctrl and *Jag1* CKO whole cochlea. The upper band (541 bp) indicates the wild-type allele containing exon 4. The lower band (286 bp, white arrow) confirms deletion of *Jag1* exon 4 in *Jag1* CKO mice. The upper band is still present in *Jag1* CKO mice since JAG1 Is expressed in all SCs and *Fgfr3-iCreER^T2/+^* only targets PCs and DCs. **(C-D)** Representative confocal slice images of HC, labeled with myosin VIIa (blue) in P7 Ctrl (C) and *Jag1* CKO (D) mice. **(E-F’)** Representative confocal slice images of SC nuclear layers at the level of HeC nuclei (E, F) and at the level of PC/DC nuclei (E’, F’) showing SOX2 (a SC marker, green) and S100a1 (a PC and DC marker, white) expression in P7 Ctrl (E-E’) and *Jag1* CKO (F-F’) mice. Arrows indicate the location of HeCs (E, F). **(G-H”)** Representative confocal slice images of SOX2 (green), FABP7 (a HeC marker, magenta), and CD44 (a CC marker, red) expression in P7 Ctrl (G-G”) and *Jag1* CKO (H-H”) mice. Arrows indicate the location of HeCs (G’, H’). **(I)** Quantification of IHCs, OHCs, and individual SC subtypes (IPCs to HeCs) in Ctrl and *Jag1* CKO mice. Data are expressed as mean ± SEM. n=4/group; *p<0.05, ****p<0.0001. Abbreviations: IHC, inner hair cell; OHC, outer hair cell; IPC, inner pillar cell; OPC, outer pillar cell; DC1, Deiters’ cell row 1; DC2, Deiters’ cell row 2; DC3, Deiters’ cell row 3; HeC, Hensen’s cell. Scale bar: 20 μm.

We next examined *Fgfr3-iCreER^T2/+^::Jag1^fx/fx^* mice at older ages to investigate whether SC loss progressed to other cell types. After tamoxifen treatment at P0 and P1, the number of HCs and individual SC subtypes were quantified in *Fgfr3-iCreER^T2/+^::Jag1^fx/fx^* cochlea and iCre-negative controls at P30 and P60 (Fig. 6A). Similar to the changes we observed at P7, there was no difference in number of IHCs or OHCs at P30 or P60 compared to the control samples (Fig. 6B-C). For SCs, the only significant difference observed at P30 and P60 was a decrease in HeCs. Thus, neonatal *Jag1* deletion induces HeC loss that occurs by P7 and does not affect other cell types or cause further morphological changes in adulthood.

**Figure 6.**
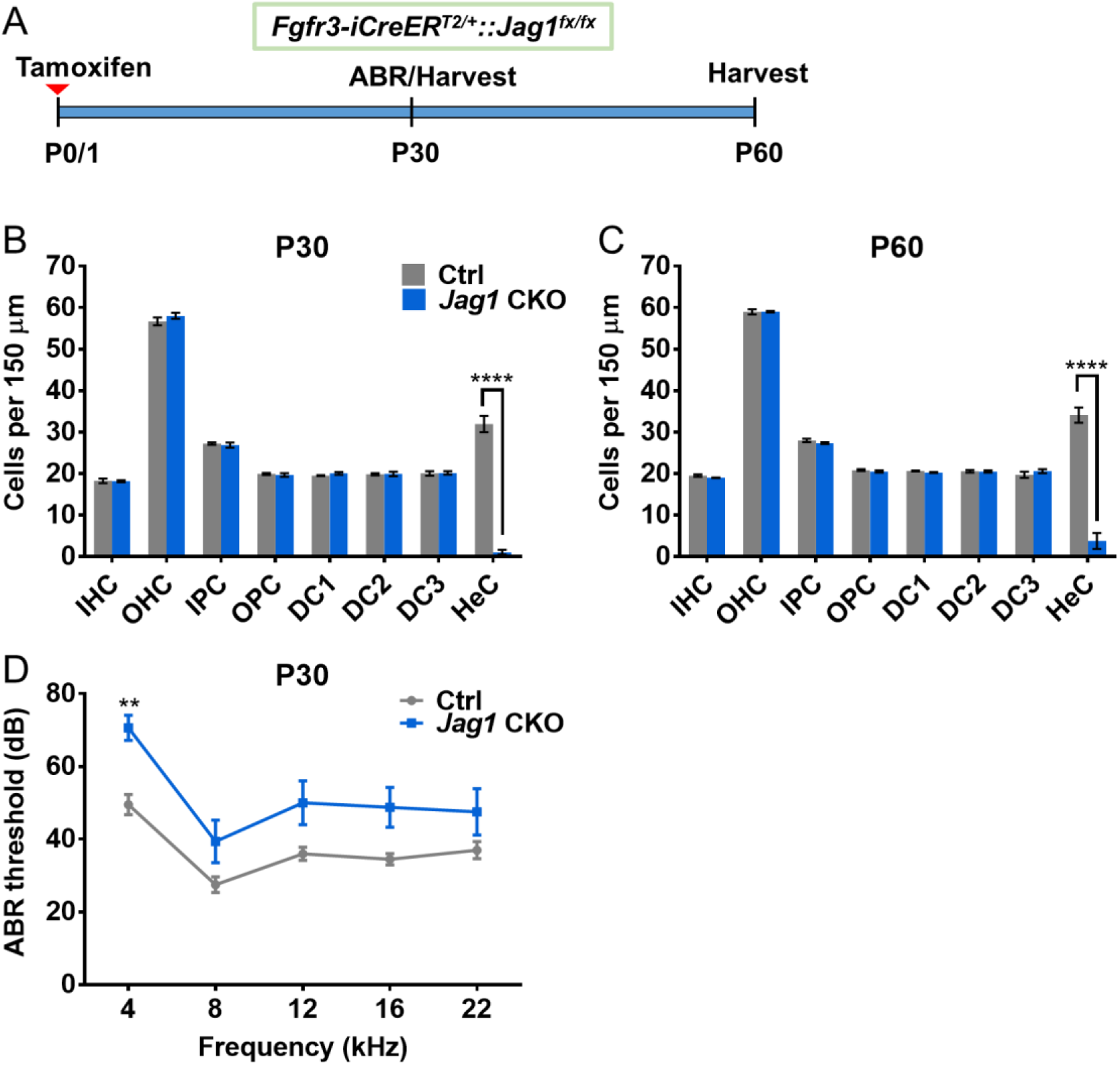
Neonatal *Jag1* deletion from PCs and DCs caused hearing deficits in the low frequency range. **(A)** Experimental strategy. *Fgfr3-iCreER^T2/+^::Jag1^fx/fx^* (*Jag1* CKO) and *Jag1^fx/fx^* (Ctrl) mice were injected with tamoxifen at P0/P1, followed by ABR at P30, and analysis of the cochlea at P30 or P60. **(B-C)** Quantification of IHCs, OHCs, and individual SC subtypes (IPCs to HeCs) in Ctrl and *Jag1* CKO mice at P30 (B) and P60 (C). Data are expressed as mean ± SEM. n=4/group; ****p<0.0001. **(D)** ABR thresholds of Ctrl and *Jag1* CKO at P30. Data are expressed as mean ± SEM. n=8-10/group; **p<0.01. Abbreviations: IHC, inner hair cell; OHC, outer hair cell; IPC, inner pillar cell; OPC, outer pillar cell; DC1, Deiters’ cell row 1; DC2, Deiters’ cell row 2; DC3, Deiters’ cell row 3; HeC, Hensen’s cell.

To assess whether HeC loss affected hearing, we performed auditory brainstem response (ABR) at P30 using *Fgfr3-iCreER^T2/+^::Jag1^fx/fx^* mice and iCre-negative controls that were injected with tamoxifen at P0 and P1 (Fig. 6A). While there was a main effect of genotype, the only significant increase in ABR thresholds in *Fgfr3-iCreER^T2/+^::Jag1^fx/fx^* mice was observed at 4 kHz (Fig. 6D).

To investigate the function of *Jag1* at an older age when HCs and SCs are more mature, we injected *Fgfr3-iCreER^T2/+^::Jag1^fx/fx^* mice with tamoxifen at P6 and P7 and quantified HCs and individual SC subtypes one week later, at P13 (Fig. 7A). Again, the numbers of IHCs and OHCs remained unchanged (Fig. 7B-D). However, unlike the neonatal *Jag1* deletion that resulted in massive HeC loss across all three cochlear turns, deletion of *Jag1* at P6/P7 induced a mild loss of HeCs that exhibited a gradient across cochlear turns. In the apical turn of *Fgfr3-iCreER^T2/+^::Jag1^fx/fx^* mouse cochleae, no HeC loss was observed (Fig. 7B). In the middle turn, there was a ~20% loss of HeCs (Fig. 7C) and in the basal turn, *Jag1* deletion at P6/P7 caused a larger ~40% loss of HeCs (Fig. 7D). Thus, there appears to be a critical period for *Jag1’*s function in SCs.

**Figure 7.**
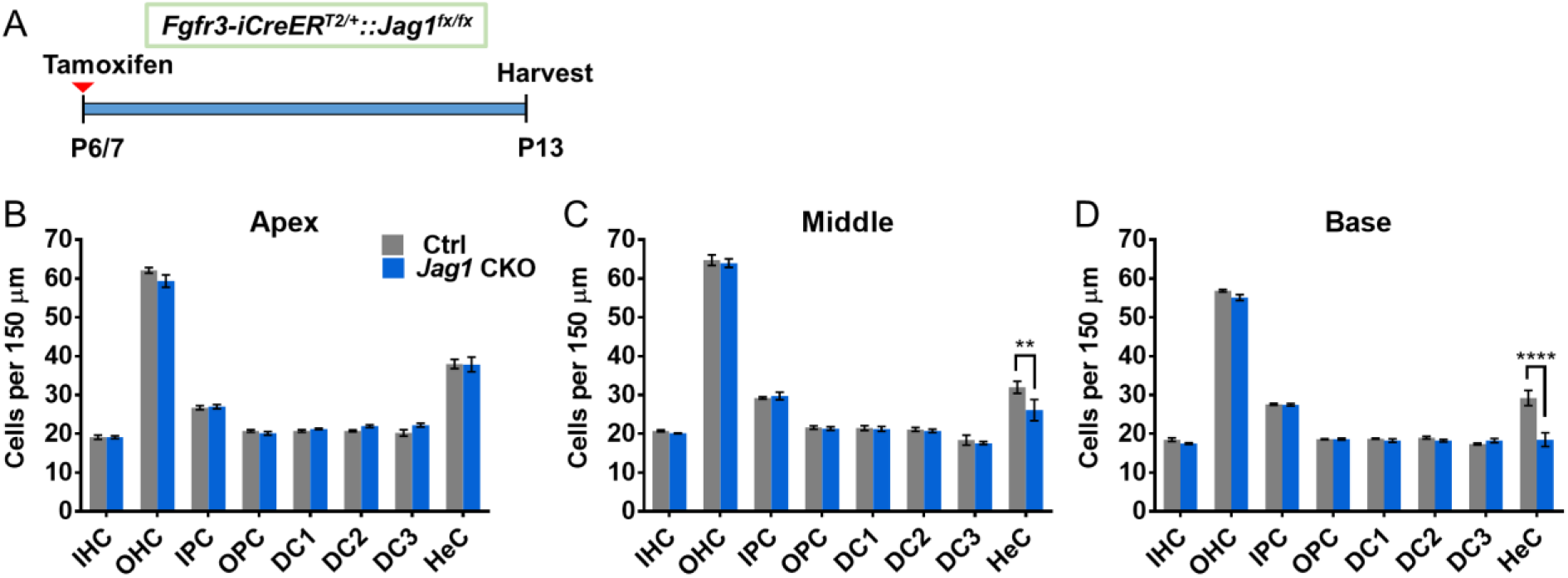
Mild loss of HeCs after deletion of *Jag1* from PCs and DCs at one week of age. **(A)** Experimental strategy. *Fgfr3-iCreER^T2/+^::Jag1^fx/fx^* (*Jag1* CKO) and *Jag1^fx/fx^* (Ctrl) mice were injected with tamoxifen at P6/P7 and cochleae are analyzed at P13. **(B-D)** Quantification of IHCs, OHCs, and individual SC subtypes (IPCs to HeCs) in the apical (B), middle (C), and basal (D) turns of the cochleae in Ctrl and *Jag1* CKO mice at P13. Data are expressed as mean ± SEM. n=4/group; **p<0.01, ****p<0.0001. Abbreviations: IHC, inner hair cell; OHC, outer hair cell; IPC, inner pillar cell; OPC, outer pillar cell; DC1, Deiters’ cell row 1; DC2, Deiters’ cell row 2; DC3, Deiters’ cell row 3; HeC, Hensen’s cell.

## DISCUSSION

JAG1 has a well-established role in specifying the prosensory region during development of the inner ear via Notch-mediated lateral induction (Eddison et al., 2000, Brooker et al., 2006, Kiernan et al., 2006, Daudet and Lewis, 2005, Hartman et al., 2010, Neves et al., 2011). However, its expression is maintained in SCs (Murata et al., 2006, Hartman et al., 2007, Oesterle et al., 2008, Maass et al., 2015) without a defined function. Here, we investigated the function of JAG1 in differentiating and maturing cochlea using two complementary *Jag1* CKO mouse models, *Sox2^CreERT2/+^::Jag1^fx/fx^* and *Fgfr3-iCreER^T2/+^::Jag1^fx/fx^*. We show using morphological and fate-mapping analyses that JAG1 is essential for the formation of HeCs and is required for HeC survival and cell fate maintenance during the early phase of cochlear maturation. Furthermore, global gene expression analysis identifies SC-specific genes as *Jag1* targets and reveals a potential link between JAG1-Notch signaling and tissue homeostasis and metabolism. Finally, functional analysis reveals that JAG1 function in the maturing cochlea is critical for low frequency hearing.

Acute disruption of Notch signaling during cochlear differentiation and early maturation triggers the conversion of SC precursors and newly formed SCs into HCs (Yamamoto et al., 2006, Tang et al., 2006, Takebayashi et al., 2007, Doetzlhofer et al., 2009). Thus, it could be reasoned that HeCs are lost due to their conversion into neighboring OHCs. However, our analyses did not reveal any increase in the number of IHCs or OHCs in response to *Jag1* deletion, indicating that JAG1 is dispensable for HC fate repression and that the observed loss of HeCs was not due to their conversion into OHCs.

While Notch signaling is most commonly known for its roles in regulating cell fate and proliferation, recent studies showed that the developing cochlear prosensory cells and later SCs, in particular DCs, depend on Notch signaling for their survival (Yamamoto et al., 2011, Basch et al., 2011, Campbell et al., 2016). Indeed, our analysis uncovered morphological evidence of dead and dying HeCs in P6/7 *Jag1* mutant cochlear tissue and our fate-mapping experiments indicate that approximately one third of HeCs are lost due to cell death. There are several studies demonstrating that Notch signaling promotes the survival of neural stem/progenitor cells (Oishi et al., 2004, Androutsellis-Theotokis et al., 2006, Mason et al., 2006) and tumor cells (Bellavia et al., 2000, Purow et al., 2005). Mechanistic work suggests that Notch activation, through non-canonical pathway(s), activates AKT, NF-κB and mTOR dependent signaling cascades, increasing oxidative metabolism and the expression of pro-survival genes (Oishi et al., 2004, Chan et al., 2007, Perumalsamy et al., 2010, Hossain et al., 2018). Consistent with this link, our gene expression analysis revealed that loss of *Jag1* resulted in a broad down-regulation of genes associated with oxidative phosphorylation and protein synthesis. Future studies will test the possible role of JAG1 in SC bioenergetics and explore possible links between JAG1-Notch signaling and pro-survival pathways such as AKT-mTOR signaling (Saxton and Sabatini, 2017).

Our fate-mapping experiments also indicate that in response to neonatal *Jag1* deletion, about two thirds of HeCs convert into CD44+, SOX2-CC-like cells. What triggers the up-regulation of CD44 and down-regulation of SOX2 in a subset of HeCs is unclear. JAG1’s role in repressing a HeC-to-CC fate switch is reminiscent of its early role in vestibular prosensory cell maintenance. There, JAG1-Notch signaling prevents the conversion of prosensory progenitors into non-sensory cells through maintaining the expression of the transcription factor SOX2 and through antagonizing the transcription factor LMX1A (Mann et al., 2017, Brown et al., 2020). A similar antagonistic relationship between CC-specific determination factors and JAG1-SOX2 axis may be at play and future investigations into the molecular mechanisms of CC fate determination are warranted.

HeCs are a SC subtype located between the 3rd row of DCs and CCs, with no direct contact with HCs. Thus, HeCs may be uniquely vulnerable to *Jag1* deletion since they only receive Notch signals from neighboring DCs through the JAG1 ligand. By contrast the Notch receptors located on DCs and PCs are also activated by the Notch ligands DLL1 and JAG2, expressed by HCs (Lanford et al., 1999; Kiernan et al., 2005; Brooker et al., 2006). Thus *Jag1* deletion using *Sox2^CreERT2/+^* or *Fgfr3-iCreER^T2^* removes all Notch signaling from HeCs; whereas DCs and PCs would still receive Notch signaling activated by DLL1 and JAG2. However, HeCs’ dependence on JAG1 appears to have a critical period as *Jag1* deletion at P6 produced a less severe phenotype where only 20-40% of HeCs were lost in the middle and basal turns of the cochlea. The milder HeC phenotype at P6 may be due to a lower Cre-mediated recombination efficiency and/or an increase in JAG1 protein stability compared to P0. The later may explain why the apex, which expresses JAG1 at a 2-fold higher levels than other regions of the cochlea, shows no loss of HeCs (Son et al., 2012).

*Jag1* deletion at the onset of cochlear differentiation not only resulted in a loss of HeCs but caused mild patterning and laminar structure defects within the auditory sensory epithelium. A recent study proposed that HeCs are involved in the patterning of OHCs during cochlear differentiation. The authors found that HeC division and migration lateral to OHCs produces a shearing motion that facilitates the arrangement of OHCs and DCs into the final checkerboard-like pattern (Cohen et al., 2019). Thus, the mild HC and SC patterning defects we observed may be the result of missing HeCs. Alternatively, the mild defects in cellular patterning in *Jag1* mutant sensory epithelia may be due to altered cell adhesion and/or polarity properties of SCs and HCs.

We also observed a decrease in the number of DCs, IPCs, and OPCs when *Jag1* was deleted at E14.5 using *Sox2^CreERT2/+^* mice compared to controls that have normal *Sox2* expression, which suggests that loss of JAG1 combined with *Sox2* haploinsufficiency affects DC and PC differentiation. Surprisingly, deletion of *Jag1* at P0 using *Sox2^CreERT2/+^* mice resulted in an opposite IPC phenotype, with a mild but significant increase in the number of IPCs compared to untreated *Sox2^CreERT2/+^::Jag1^fx/fx^* controls but not compared to controls that have normal *Sox2* expression. These results emphasize the need for controls of various genotypes and the use of different methods to study gene deletions since genes may interact in networks.

We found that *Jag1* deletion at P0 using *Fgfr3-iCreER^T2^* resulted in mild but significant hearing deficits at 4 kHz. The reason for the decrease in low frequency hearing sensitivity is unknown. However, it is intriguing to speculate that it is causally linked to diminished HeC function. HeCs have a role in the formation of the tectorial membrane (Rau et al., 1999, Kim et al., 2019), they participate in the recycling of potassium ions from the base of OHCs to the endolymph through gap junction connections with DCs (Sato and Santos-Sacchi, 1994, Kikuchi et al., 1995, Zwislocki et al., 1992, Lautermann et al., 1998), and they are involved in maintaining osmotic balance through their expression of aquaporin4 (Takumi et al., 1998, Li and Verkman, 2001). Moreover, a recent study provides evidence that the passive mechanical properties of HeCs may help tune the OHC force generation in the cochlear apex (Gao et al., 2014). Deregulation of any of these processes could alter hearing sensitivity, with the later altering high frequency sensitivity.

While HeCs were discovered over a century ago, little was known about the factors and signals that control their development. In this study, we identify the Notch ligand JAG1 as being essential for HeC differentiation and maturation, revealing a novel role for JAG1 in SC-fate maintenance and SC survival.

## MATERIALS AND METHODS

### Animals

*Sox2^CreERT2/+^* mice (Arnold et al., 2011) (stock #17593) and *Rosa26^loxP-stop-loxP-tdTomato^* (*Rosa26^tdTomato^*) mice (Madisen et al., 2010) (stock #7914) were purchased from The Jackson Laboratory (Bar Harbor, ME). *Fgfr3-iCreER^T2^* mice (Rivers et al., 2008, Young et al., 2010) were provided by Dr. William Richardson (University College, London). Two different *Jag1^fx/fx^* mouse lines were used in this study. For *Jag1* deletion using *Sox2^CreERT2/+^* mice, *Jag1^fx/fx^* mice (Brooker et al., 2006) (MGI: 3623344) were provided by Dr. Julian Lewis (Cancer Research UK London Research Institute). In this line, loxP sites were inserted to flank exons 4 and 5, resulting in a frame shift that generates a stop codon after the start of exon 6 following Cre-mediated recombination. For *Jag1* deletion using *Fgfr3-iCreER^T2/+^* mice and fate-mapping with *Sox2^CreERT2/+^* mice, *Jag1^fx/fx^* mice (Kiernan et al., 2006) (MGI: 3692444) (stock# 10618) were purchased from The Jackson Laboratory (Bar Harbor, ME). In this line, loxP sites were inserted to flank exon 4, resulting in a non-functional JAG1 protein after Cre-mediated recombination. Genotyping was performed in house using PCR as previously described for each line or by Transnetyx, Inc. (Cordova, TN). Mice of both genders were used in all studies. All animal work was performed in accordance with approved animal protocols from the Institutional Animal Care and Use Committees at Johns Hopkins University or Southern Illinois University School of Medicine.

### Substances given to animals

*Sox2^CreERT2/+^::Jag1^fx/fx^* mice: Pregnant dams time-mated to E14.5, received two intraperitoneal (IP) injections of tamoxifen (0.125 mg/g body weight, cat #T5648, Sigma-Aldrich, St. Louis, MO) and progesterone (0.125 mg/g body weight, cat #P3972 Sigma-Aldrich, St. Louis, MO), given 8 hours apart. Neonatal pups were injected with (Z)-4-hydroxytamoxifen (0.139 mg per pup, cat #H7904-5mg, Sigma-Aldrich, St. Louis, MO) at P0 and P1. Controls included Cre-negative littermates, as well as *Sox2^CreERT2/+^*::*Jag1^fx/fx^* and *Sox2^CreERT2/+^* mice that did not receive tamoxifen to control for *Sox2* haploinsufficiency. *Fgfr3-iCreER^T2/+^::Jag1^fx/fx^* mice and *Sox2^CreERT2/+^:: Jag1^fx/fx^::Rosa26^tdTomato^* mice were injected with tamoxifen (0.075mg/g body weight, cat #T5648, Sigma-Aldrich, St. Louis, MO) at P0 and P1, or at P6 and P7. Controls were iCre-negative littermates or *Sox2^CreERT2/+^::Rosa26^tdTomato^* littermates that also received tamoxifen.

### Tissue processing

Late embryonic and early postnatal animals were staged using the EMAP eMouse Atlas Project (http://www.emouseatlas.org) Theiler staging criteria. Temporal bones were collected and fixed in 4% paraformaldehyde (PFA, Polysciences, Inc., Warrington, PA) in 10 mM PBS for 2 hours at room temperature. Cochleae were then dissected using the whole mount method as previously described (Montgomery and Cox, 2016). For cryo-sectioning, PFA-fixed inner ears were put through a sucrose gradient (10% sucrose for 30 minutes, 15% sucrose for 30 minutes, and 30% sucrose overnight), submerged in OCT (Tissue-Tek, Sakura Finetek USA, Torrance, CA) and flash frozen. Sections cut at 14 μm thickness were collected on SuperFrost Plus slides (Fisher, Hampton, NH).

### Immuno-staining

Cochlear samples were immuno-stained following standard procedures (Montgomery and Cox, 2016). Briefly, tissue was blocked and permeabilized in 10 mM PBS containing 10% Normal Horse Serum (NHS) or Normal Goat Serum (NGS) (Vector Labs, Burlingame, CA), 1% Bovine Serum Albumin (BSA) (Fisher, Hampton, NH), and 1% Triton-X-100 (Sigma-Aldrich, St. Louis, MO) for 1 hour at room temperature. Then samples were incubated with primary antibodies diluted in 10 mM PBS containing NHS or NGS (5%), BSA (1%), and Triton-X-100 (0.1%) overnight at 4°C or 37°C. The following primary antibodies were used: rabbit anti-FABP7 (1:200, cat #ab32423, Abcam, Cambridge, MA), rat anti-CD44 (1:500, cat #550538, BD Pharmingen, San Jose, CA), rabbit anti-cleaved caspase3 (1:1000, cat #9661, Cell Signaling Technology, Danvers, MA), goat anti-JAG1 (1:500, cat #sc-6011, Santa Cruz Biotechnology, Dallas, TX), rabbit anti-myosin VIIa (1:200, cat #25-6790, Proteus BioSciences, Ramona, CA), mouse anti-parvalbumin (1:1000, cat #P3088, Sigma-Aldrich, St. Louis, MO), rabbit anti-S100a1 (1:400, cat #ab868, Abcam, Cambridge, MA), and goat anti-SOX2 (1:400, cat #sc-17320, Santa Cruz Biotechnology, Dallas, TX). The next day, samples were incubated in corresponding Alexa Fluor-conjugated secondary antibodies (1:1000, Life Technologies, Eugene, OR) for 2 hours at room temperature. Nuclei were stained with Hoechst 33342 (1:2000, cat #H1399, Fisher, Hampton, NH) for 20 minutes at room temperature, followed by mounting on slides in Prolong Gold (cat #P36930, Fisher, Hampton, NH). To increase the signal to noise ratio, some samples stained with anti-SOX2 antibodies were pretreated with signal enhancer (cat #136933, Life Technologies, Eugene, OR) for 30 min at room temperature before the blocking and permeabilization step. Samples were imaged using a Zeiss LSM 800 confocal microscope and processed with Zen Blue and ZEN 2.3 SP1 software (Carl Zeiss, Oberkochen, Germany).

### Cell counts

Confocal z-stacks through the HC layer and corresponding SC layer were taken at two representative regions in each turn of the cochlear (apex, middle, and base). Labeled IHCs, OHCs, and different SC subtypes were quantified in two 150 or 200 μm representative regions in each turn of the cochlea. The length of the imaged segment (150 or 250 μm) was analyzed using Image J (http://imagej.nih.gov/ij) or Zen Blue software. HCs and SCs were manually counted in Photoshop CS5 (Adobe, San Jose, CA) or in Zen Blue software.

### Validation of *Jag1* deletion

Immuno-staining with anti-JAG1 antibodies was used to validate deletion of *Jag1* from SCs using the *Jag1^fx/fx^* allele that was provided by Dr. Julian Lewis (Brooker et al., 2006). However, the *Jag1^fx/fx^* allele purchased from The Jackson Laboratories produces a truncated protein that was still detected by anti-JAG1 antibodies. Therefore, we used RT-PCR to validate the deletion of *Jag1* from SCs in these mice. Total RNA was extracted from P7 *Jag1^fx/fx^* and *Fgfr3-iCreER^T2/+^::Jag1^fx/fx^* cochleae using tri-reagent (cat #T9424, Sigma-Aldrich, St. Louis, MO). RNA was then precipitated with isopropanol (cat #327930010, Acros Organics, Geel, Belgium) overnight at −20°C, purified with DNAse treatment (cat #AM1906, Fisher, Hampton, NH) and re-precipitated with sodium acetate (cat #BP333, Fisher, Hampton, NH). Reverse transcription into cDNA was performed using Thermo Scientific Maxima First Strand cDNA synthesis kit (cat #K1641, Fisher, Hampton, NH, United States). cDNA was amplified with primers that flanked exon 4 of *Jag1* [forward: CGA CCG TAA TCG CAT CGT AC, reverse: AGT CCC ACA GTA ATT CAG ATC; (Kiernan et al., 2006)], separated on a 1% agarose gel and visualized with GelRed nucleic acid stain (cat #41001, Biotium, Fremont, CA) using a Syngene G:Box Chemi imager (Syngene, Frederick, MD).

### Microarray and RT-qPCR experiments

For microarray and qPCR validation experiments control (*Jag1^fx/fx^*) and experimental animals (*Sox2^CreERT2/+^::Jag1^fx/fx^*) were harvested at P0 after receiving tamoxifen and progesterone at stage E14.5. Cochlear epithelia from individual animals were isolated using dispase/collagenase digest as previously described (Doetzlhofer et al., 2009). Total RNA was extracted using the RNeasy Micro Kit (cat #74004, QIAGEN, Germantown, MD). For the microarray experiment, total RNA was labeled with the Affymetrix GeneChip WT PLUS Reagent Kit using the manufacturer’s protocol. Labeled RNA was hybridized onto Clariom D mouse microarray (Affymetrix, Santa Clara, CA) and chips were scanned and analyzed according to manufacturer’s manuals. Affymetrix CEL files were extracted and data normalized with the Partek GS 6.6 platform (Partek Inc., Chesterfield, MO). Partek’s extended meta-probe set was used with RMA normalization to create quantile-normalized log2 transcript signal values, which were used in subsequent ANOVA analyses. The microarray data is deposited in the Gene Expression Omnibus (GEO) data base, accession number GSE148009. Results were validated using qPCR experiments where mRNA was reverse transcribed into cDNA using the iScript cDNA synthesis kit (cat #1708891, Bio-Rad, Hercules, CA). SYBR Green based qPCR was performed on a CFX-Connect Real Time PCR Detection System (Bio-Rad, Hercules, CA) using Fast SYBR® Green Master Mix reagent (cat #4385616, Applied Biosystems, Foster City, CA) and gene-specific primers. Relative gene expression was analyzed using the comparative CT method (Schmittgen and Livak, 2008). The ribosomal gene *Rpl19* was used as reference gene and wild-type early postnatal cochlear tissue was used as calibrator. The following qPCR primers were used:

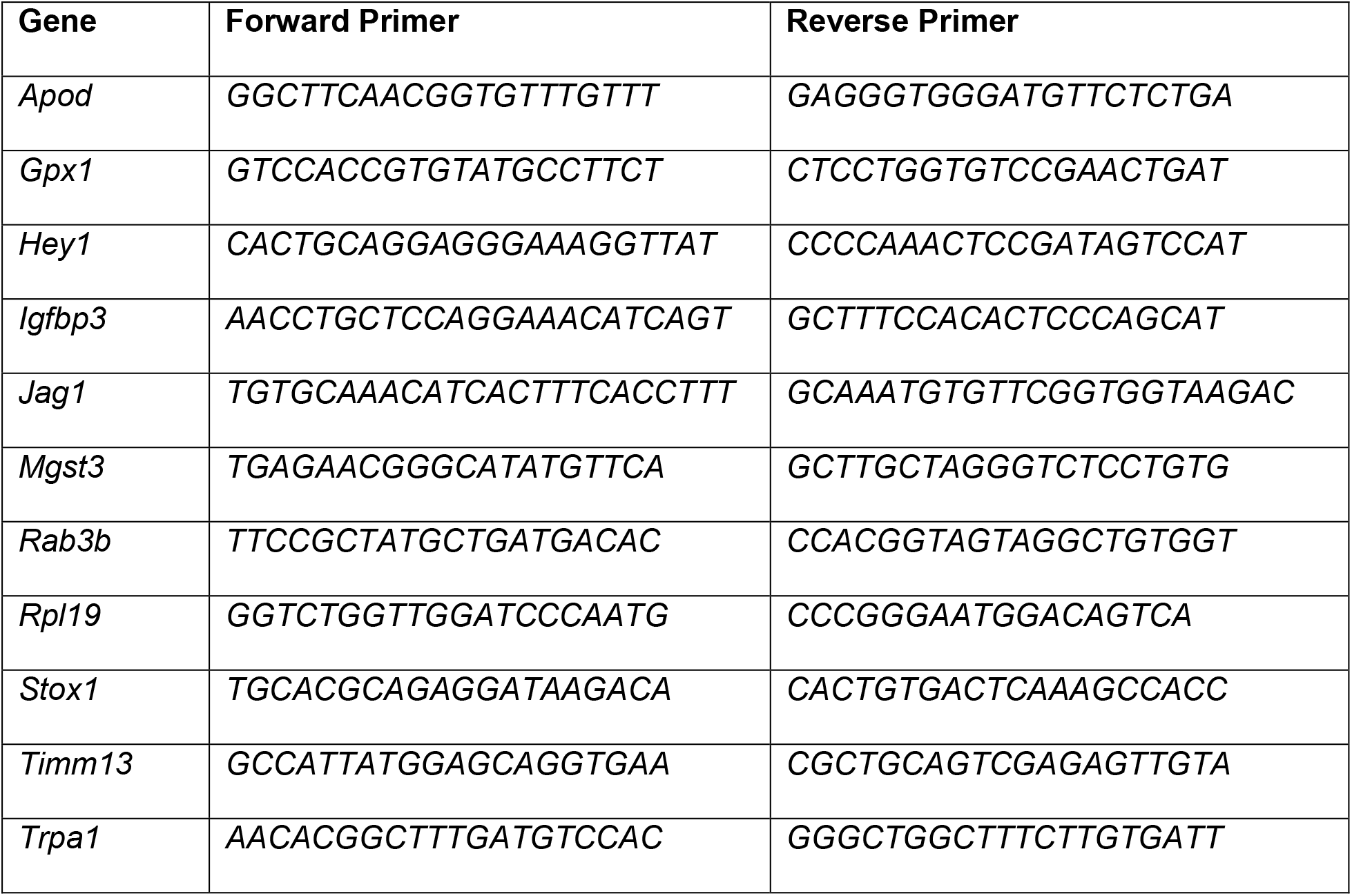

### Gene Ontology Analysis

Gene ontology (GO) enrichment analysis was conducted using the Functional Annotation Tool in DAVID (Huang da et al., 2009b, Huang da et al., 2009a). A curated list of down-regulated genes (p-value ≤0.05, log2 (FC) ≤ −2σ, mean log2 Ctrl ≥5) was uploaded into the DAVID bioinformatics suite 6.8 and GO terms related to biological process (GOTERM BP DIRECT), molecular function (GOTERM MF DIRECT), cellular compartment (GOTERM CC DIRECT), and KEGG pathway term (KEGG PATHWAY) were obtained using default parameters.

### Auditory brainstem response (ABR) measurements

To assess hearing after *Jag1* deletion, ABR was measured at P30 in *Fgfr3-iCreER^T2/+^::Jag1^fx/fx^* mice and iCre-negative littermates injected with tamoxifen at P0/P1. Before ABR recordings, mice were anesthetized with Avertin (250-500mg/kg, IP, Sigma-Aldrich, St. Louis, MO), and kept on a heating pad in a sound attenuated booth (Industrial Acoustic, Bronx, NY) throughout the whole procedure. One subdermal stainless-steel recording electrode was inserted at the vertex of the skull, with a second reference electrode inserted under the pinna of the left ear. A ground electrode was inserted at the base of the tail. An electrostatic speaker (EC1, Tucker Davis Technologies [TDT] System III, TDT, Alachua, FL) was fitted to a tube and inserted into the left ear canal. Acoustic signals were generated using a 16-bit D/A converter (RX6, TDT System III, TDT, Alachua, FL), controlled by a customized Auditory Neurophysiology Experiment Control Software (ANECS, Blue Hills Scientific, Boston, MA). To generate ABRs, pure tone bursts at 4, 8, 12, 16, and 22 kHz were presented at a rate of 19/sec, with a 5 ms duration, and repeated 512 times. Sound intensity at each frequency decreased from 80 to 5 dB, with 5 dB steps. Two blinded reviewers determined the ABR thresholds, which were the lowest sound intensity that produced a visually distinct response in wave I or II.

### Statistical Analysis

The sample size (n) represents the number of animals analyzed per group. Animals (biological replicates) were allocated into control or experimental groups based on genotype and/or type of treatment. To avoid bias, masking was used during data analysis. P-values ≤ 0.05 were considered significant. Data was analyzed using Graphpad Prism 6.02 and 8.0 (Graphpad Software Inc., La Jolla, CA).

## ACKNOWLEDGEMENTS

We thank Dr. William Richardson at University College London for providing the *Fgfr3-iCreER^T2^* mouse line, and Dr. Julian Lewis at Cancer Research UK London Research Institute for providing one of the *Jag1^fx/fx^* mouse lines. This work was supported by grants from NIDCD [R01DC011571 (AD) and R01DC014441 (BC)] and the Office of the Assistant Secretary of Defense for Health Affairs [W81XWH-15-1-0475 (BC)]. The Southern Illinois University School of Medicine Research Imaging Facility is supported by a grant from the Office of Naval Research (N00014-15-1-2866).

## SUPPLEMENTAL MATERIAL

**Supplemental Table 1.**
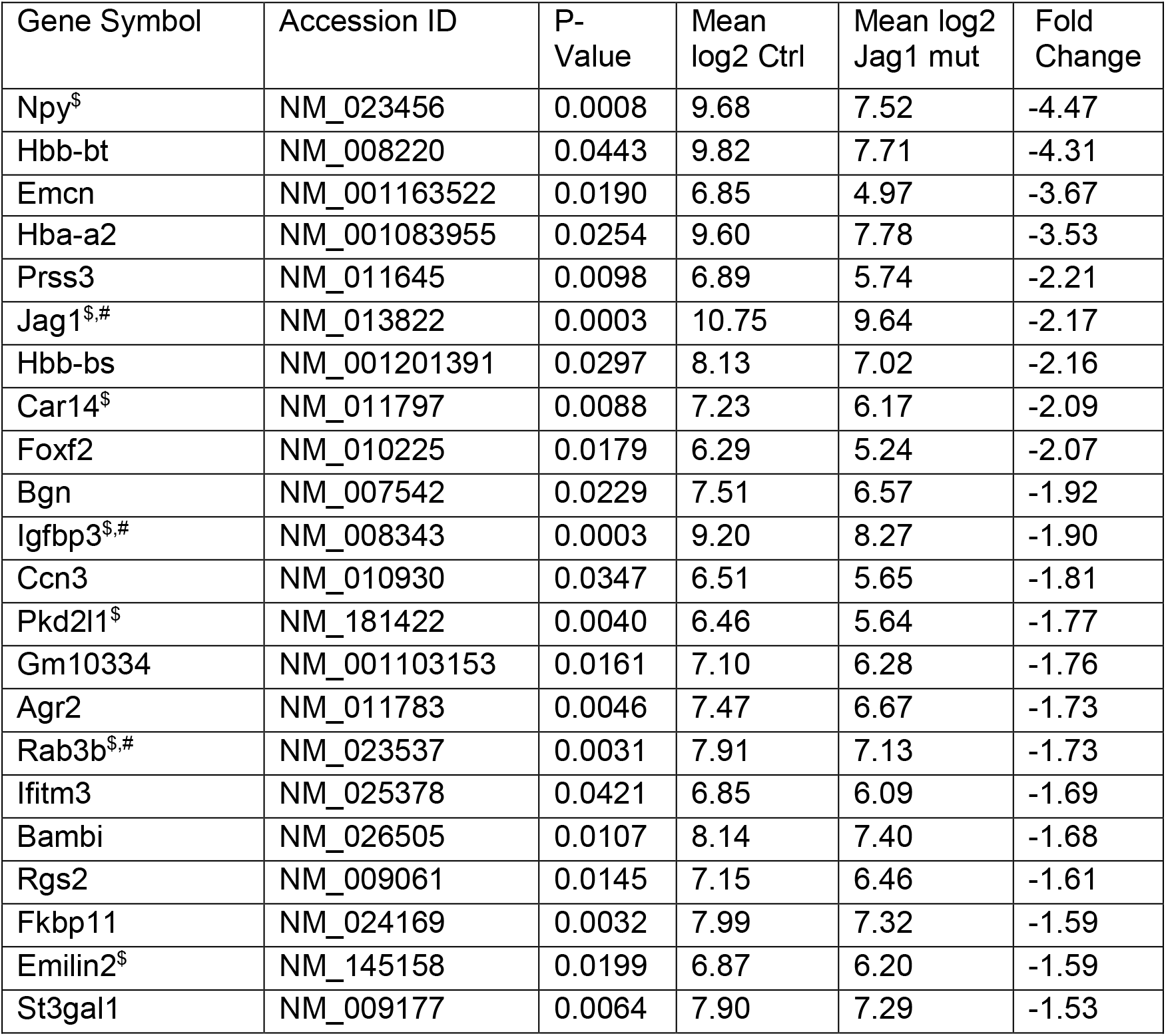
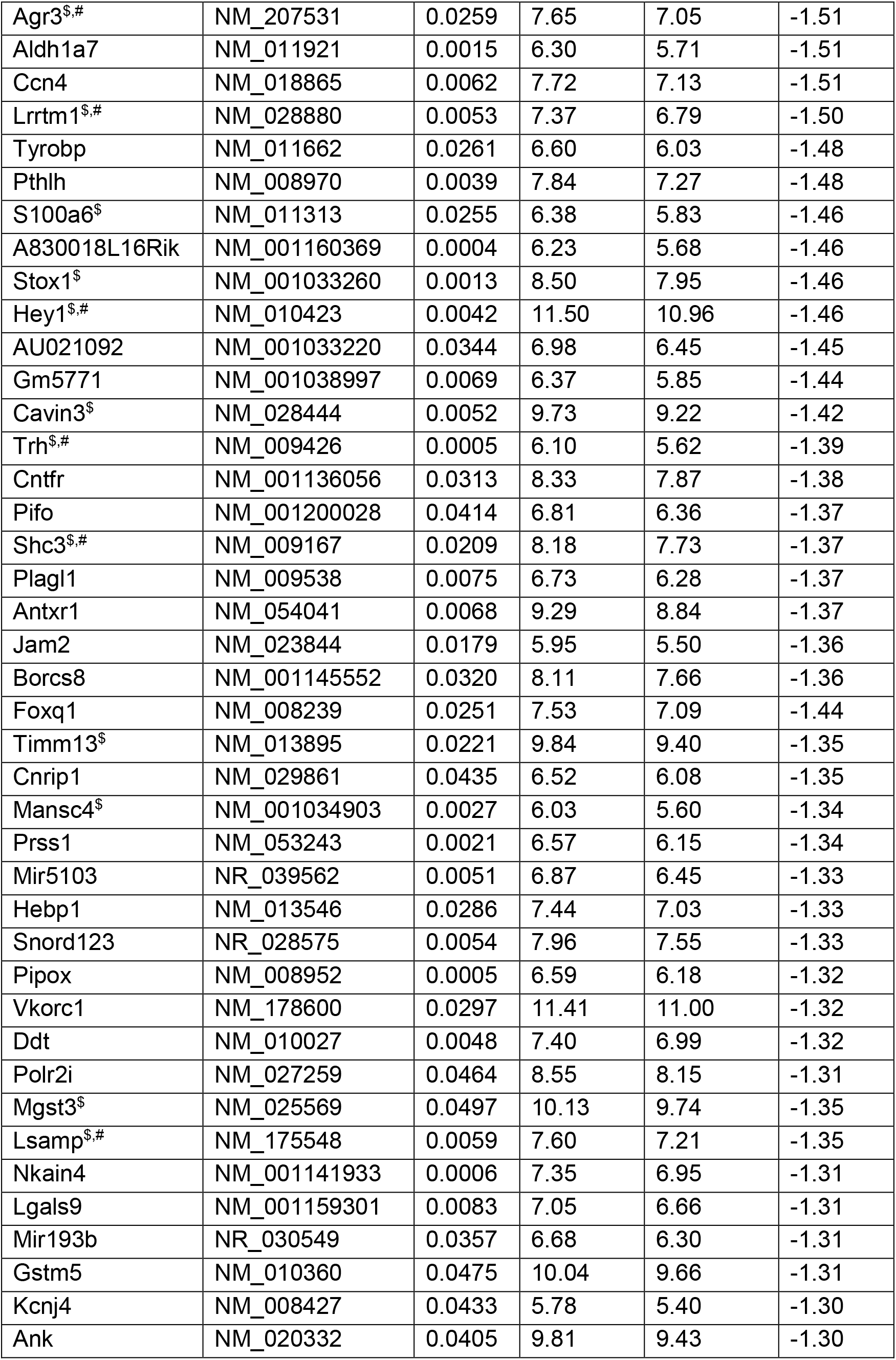
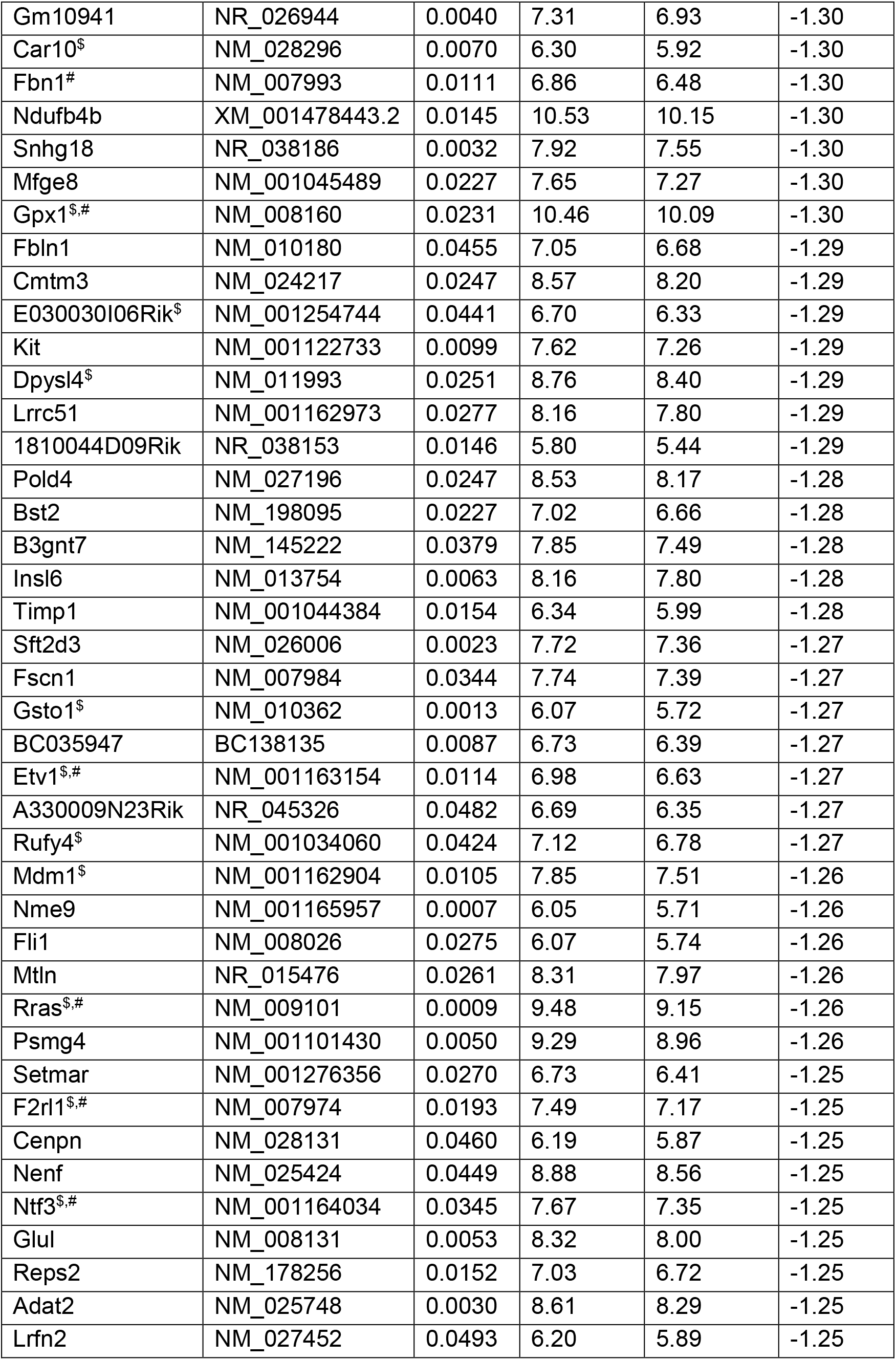
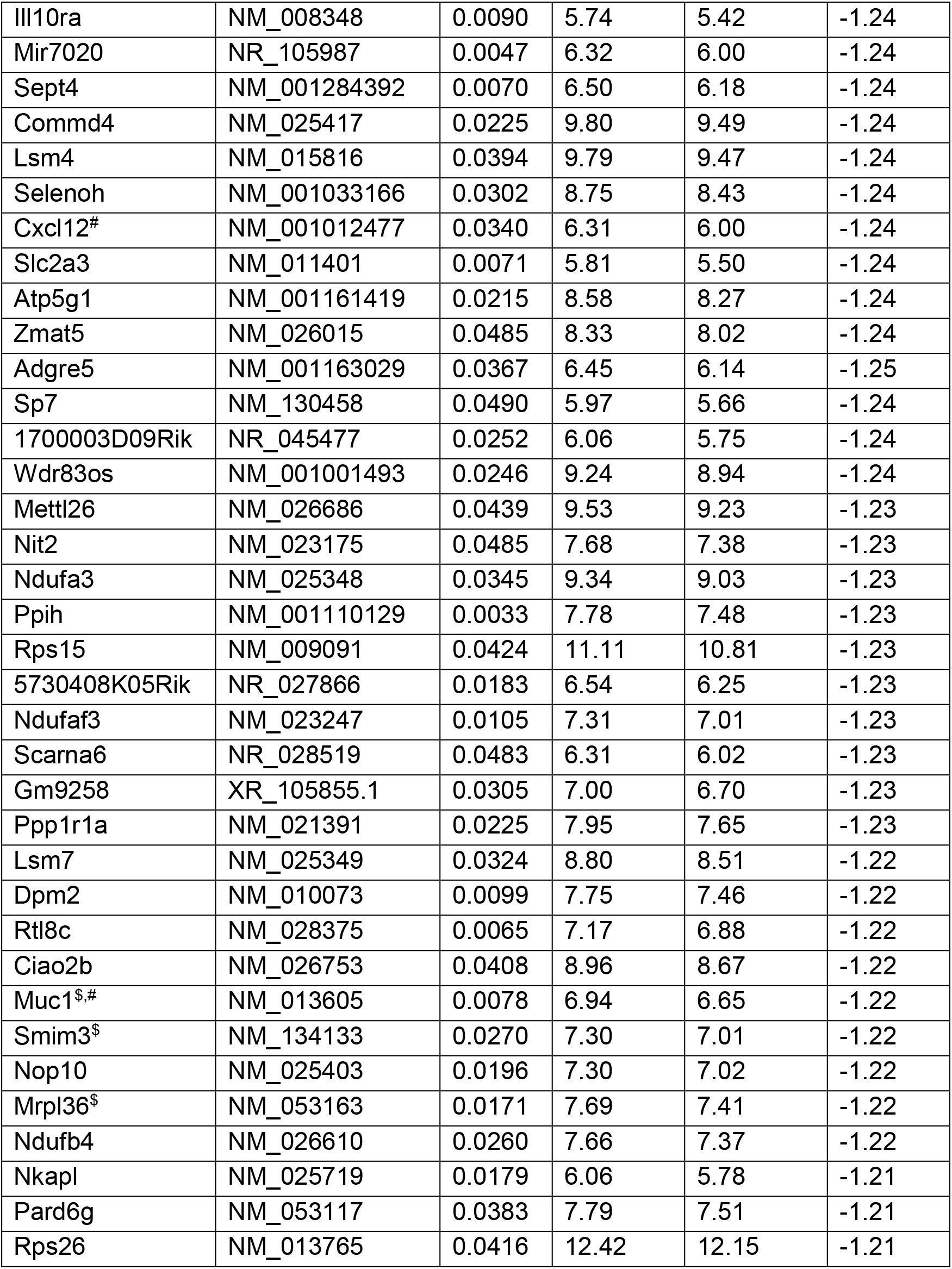
List of top-ranked down-regulated genes after *Jag1* deletion. Genes are ranked based on fold change. Shown are genes that are significantly down-regulated in *Jag1* CKO samples compared to control (p-value ≤0.05, log2 (FC) ≤3σ) after tamoxifen and progesterone injection at E14.5. $ marks SC-specific/enriched genes (Maass et al., 2016, Burns et al., 2015) and # marks genes previously found to be positively regulated in cochlear epithelial cells by Notch signaling (Maass et al., 2016, Campbell et al., 2016). Note that only transcripts with mean log2 values of 5.5 or more in control samples are listed.

**Supplemental Table 2.**
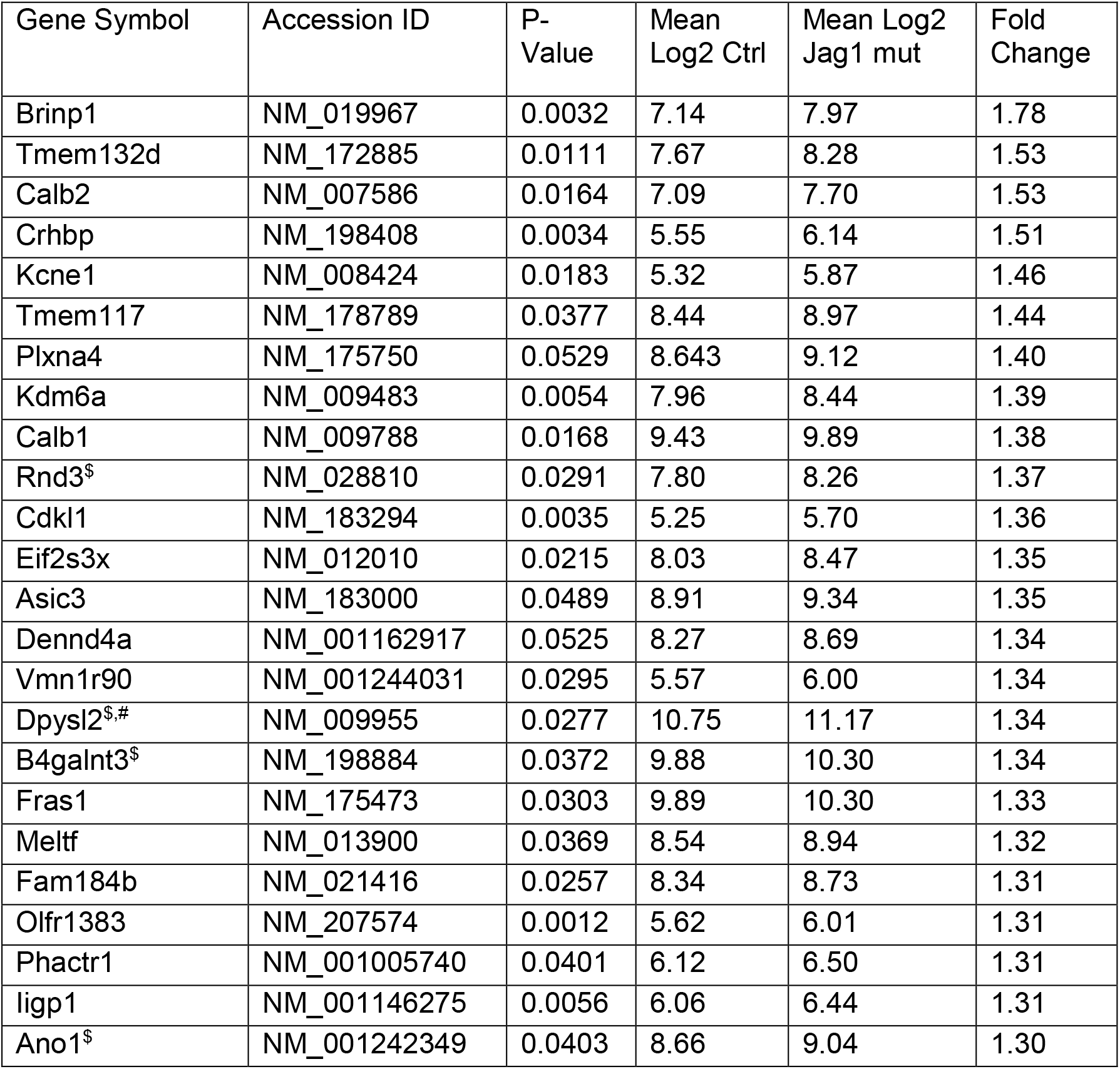
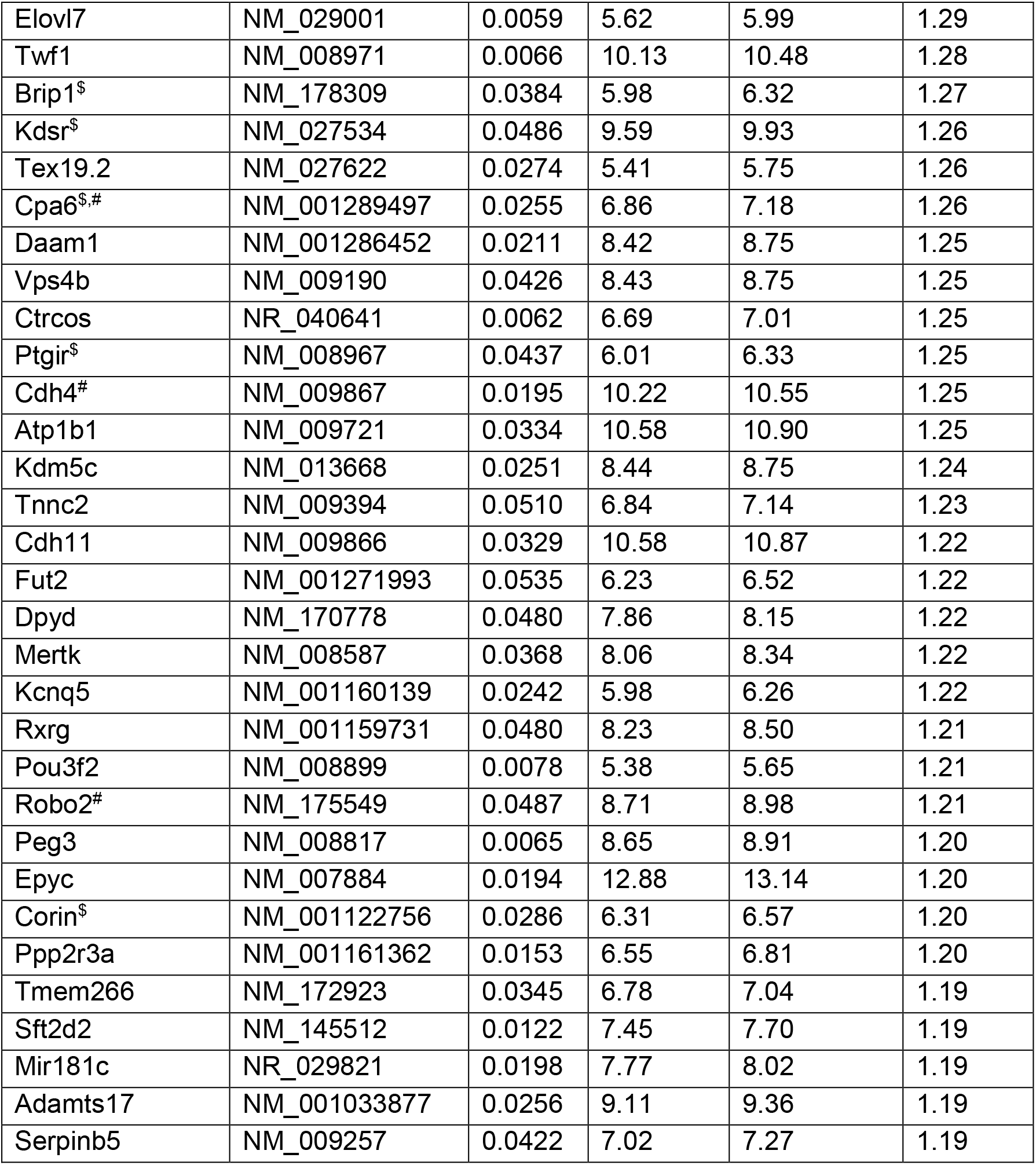
List of top-ranked up-regulated genes after *Jag1* deletion. Genes are ranked based on fold change (FC). Shown are genes that are significantly up-regulated in *Jag1* CKO samples compared to control (p-value ≤0.05, log2 (FC) ≥3σ) after tamoxifen and progesterone injection at E14.5. $ marks SC-specific/enriched genes (Maass et al., 2016, Burns et al., 2015) and # marks genes previously found to be negatively regulated in cochlear epithelial cells by Notch signaling (Maass et al., 2016, Campbell et al., 2016). Note that only transcripts with mean log2 values in Jag1 mutant samples of 5.5 or more are listed.

